# An alternative miRISC targeting a coding mutation site in *FOXL2* links to granulosa cell tumor

**DOI:** 10.1101/2020.02.18.954487

**Authors:** Eunkyoung Shin, Hanyong Jin, Dae-Shik Suh, Yongyang Luo, Hye-Jeong Ha, Tae Heon Kim, Yoonsoo Hahn, Seogang Hyun, Kangseok Lee, Jeehyeon Bae

## Abstract

Recent evidence suggests that animal microRNAs (miRNAs) can target coding sequences (CDSs); however, the pathophysiological importance of such targeting remains unknown. Here, we show that a somatic heterozygous missense mutation (c.402C>G; p.C134W) in *FOXL2*, a feature shared by virtually all adult-type granulosa cell tumors (AGCTs), introduces a target site for miR-1236, which induces haploinsufficiency of the tumor-suppressor *FOXL2*. This miR-1236-mediated selective degradation of the variant *FOXL2* mRNA is preferentially conducted by a distinct miRNA-loaded RNA-induced silencing complex (miRISC) directed by the Argonaute3 (AGO3) and DHX9 proteins. In both patients and mouse model of AGCT, the inversely regulated variant *FOXL2* abundance with the miR-1236 levels was highly correlated with malignant features of AGCT. Our study provides a molecular basis for understanding the conserved *FOXL2* CDS mutation-mediated etiology of AGCT, revealing the existence of a previously unidentified mechanism of miRNA-targeting disease-associated mutations in the CDS by forming a non-canonical miRISC.

## INTRODUCTION

MicroRNAs (miRNAs) are endogenous, noncoding RNAs of ∼22 nucleotides (nt) in length that suppress the stability or translational efficiency of mRNAs. Conventionally, miRNAs are known to target sequences in the 3′-untranslated regions (UTRs) of mRNAs. However, recent multiple high-throughput sequencing and proteomic analyses suggest that miRNAs can also bind sites within mRNA coding sequences (CDSs) (Broughton, Lovci *et al*., 2016, Chi, Zang *et al*., 2009, Hafner, Landthaler *et al*., 2010, Hausser, Syed *et al*., 2013, Schnall-Levin, Zhao *et al*., 2010, Xue, Ouyang *et al*., 2013), and the intracellular effects of miRNAs targeting CDS sites have been proposed (Elcheva, Goswami *et al*., 2009, Forman, Legesse-Miller *et al*., 2008, Schnall-Levin, Rissland *et al*., 2011, Tay, Zhang *et al*., 2008, Zhang, Zhang *et al*., 2018). However, the physiological relevance and the pathological consequences of miRNA binding to CDSs remain unclear.

Argonaute (AGO) clade proteins are essential components of miRNA-loaded RNA-induced silencing complexes (miRISCs) that select target mRNAs by directly interacting with mature miRNAs (Czech & Hannon, 2011, Hock & Meister, 2008). In mammals, four AGO paralogs (AGO1–4) are involved in miRNA pathways (Czech & Hannon, 2011, Hock & Meister, 2008), and they share ∼80% amino acid-sequence identity (Sasaki, Shiohama *et al*., 2003). AGO2 has been described as a specialized AGO that possesses slicer activity, enabling cleavage of target mRNAs by miRNAs and small-interfering RNAs (siRNAs) (Liu, Carmell *et al*., 2004, Meister, Landthaler *et al*., 2004). However, previous data suggested that all mammalian AGOs may serve overlapping and distinct functions in miRNA-mediated regulation (Su, Trombly *et al*., 2009). High-throughput pyrosequencing data showed that the profiles of miRNAs associated with AGO2 and AGO3 largely overlap, but preferential associations with AGO2 or AGO3 also occur for a small set of miRNAs (Azuma-Mukai, Oguri *et al*., 2008). AGO3 is also associated with slicer activity (Park, Phan *et al*., 2017). However, the functional significances of mammalian AGO1, AGO3, and AGO4 in miRNA activity are poorly understood.

Conclusive evidence demonstrating clear pathophysiological consequences elicited by miRNA-binding to the CDSs of disease-associated gene loci is lacking. Here, we investigated whether miRNA binding to the CDS of *FOXL2* contributes to adult-type granulosa cell tumor (AGCT) development. GCTs are malignant ovarian cancers comprised of AGCTs and juvenile GCTs (JGCTs) (Schumer & Cannistra, 2003). *FOXL2* is evolutionarily conserved and encodes a forkhead-domain transcription factor essential for the ovary development and function (Schmidt, Ovitt *et al*., 2004, Uhlenhaut, Jakob *et al*., 2009). A highly prevalent heterozygous somatic missense mutation (c.402C>G; p.C134W) in *FOXL2* is exclusively found in >97% of patients with ACGT and is considered the main cause of AGCT (Shah, Kobel *et al*., 2009). However, the etiological nature of the 402C>G mutation remains largely unknown. Previously, we showed that the FOXL2 protein acted as a tumor suppressor in granulosa cells, whereas C134W FOXL2 did not, due to serine 33 hyperphosphorylation by GSK3β, leading to accelerated MDM2-mediated ubiquitination and proteasomal degradation (Kim, Kim *et al*., 2014, Kim, Yoon *et al*., 2011). However, a relatively moderate change in FOXL2 protein stability by the C134W mutation does not appear to wholly account for haploinsufficiency of FOXL2 (Kim *et al*., 2014, Kim *et al*., 2011). Here, we identified allelic imbalance in *FOXL2* mRNAs in patients with AGCT arising from recognition of the 402C>G locus as a target site of miR-1236 that drives degradation of this variant *FOXL2* mRNA, which explains the etiology of this conserved mutation in AGCTs.

## RESULTS

### Allelic imbalance of *FOXL2* transcripts in AGCT samples

To study allelic imbalance of heterozygous *FOXL2* mRNAs, we analyzed the relative levels of wild-type (WT) and variant *FOXL2* (402C>G) mRNAs from complementary DNA (cDNA) samples from the individual AGCT tissues by high-throughput ultra-deep RNA sequencing. Ultra-deep RNA sequencing analysis of AGCT tissues showed that decreased proportion of variant *FOXL2* mRNA level compared to WT *FOXL2* mRNA in 14 AGCT patients (WT:402C>G = 73:27) and the opposite trend in 6 patients (WT:402C>G = 36:64) (Fig 1A). A previous study, which identified this conserved mutation, reported relative abundance of WT versus variant *FOXL2* mRNA levels in four AGCT patients, where no uniformed trend was observed (Shah *et al*., 2009). For these reasons, we recruited additional AGCT patients from two independent hospitals and performed high-throughput pyrosequencing analysis (n = 46) that enables amplification and detection of both alleles from the same pyrosequencing reaction using common primers designed to bind *FOXL2* cDNA. As shown in Fig 1B and Appendix Fig S1A, the relative abundance of *FOXL2* mRNA analyzed by pyrosequencing were 70:30 for WT:402C>G in 46 AGCTs including 20 corresponding AGCTs analyzed for RNA sequencing presented in Fig 1A. In addition, allele-specific real-time and semi-quantitative RT-PCR analyses of 46 AGCTs were performed using primers presented in Appendix Fig S1B, and we observed consistent lower steady-state levels of *FOXL2* variant mRNA compared to WT *FOXL2* mRNA (Appendix Fig S1C and S1D). For these analyses, we used paired genomic DNA (gDNA) levels of both alleles for the normalization of data, where the gDNA levels of both alleles were similar in all AGCTs (Appendix Fig S1E). These results indicate that contamination of non-cancerous stromal cells in preparation of total RNA from AGCT tissues for these analyses was minimal. We also performed primer extension assays on cDNA samples, as shown in Appendix Fig S1F, the mutated allele was not readily detectable in AGCTs.

**Figure 1.**
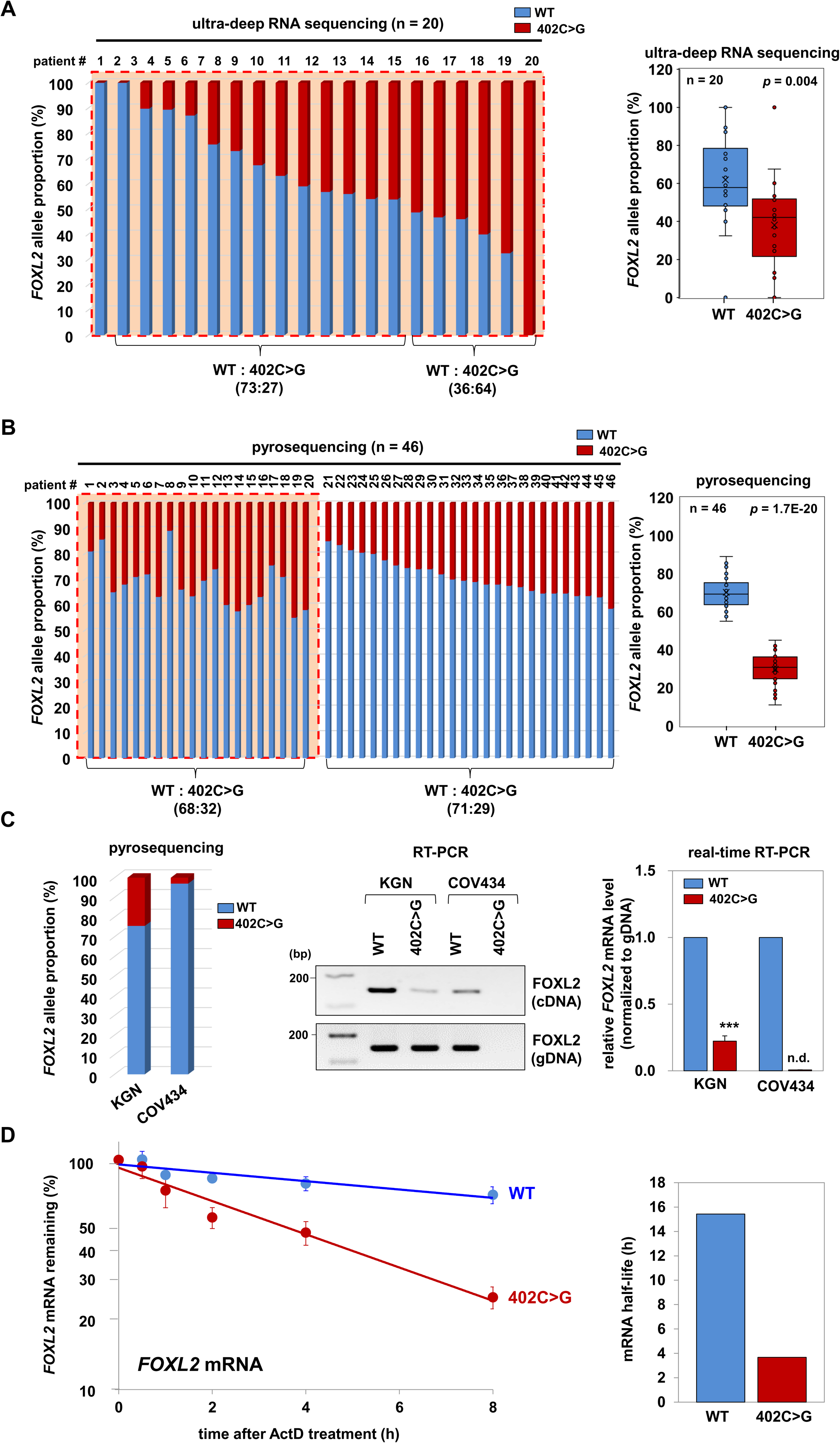
Allelic imbalance of heterozygous *FOXL2* transcripts in AGCT cells. **A, B** Bar graph and box-and-whisker plots are presented, which show the allelic proportions of WT *FOXL2* mRNA and 402C>G *FOXL2* mRNA in AGCT tissues from 20 patients analyzed by ultra-deep RNA sequencing (A) and from 46 patients analyzed by pyrosequencing (B). The box plot represents the lower, median, and upper quartiles, and the whiskers represent the 95% confidence interval of the mean. The whiskers extend to the most extreme data points not considered outliers, and the outliers are represented as dots. Orange dotted boxes indicate that AGCTs from 20 patients analyzed for both ultra-deep RNA sequencing and pyrosequencing. Comparisons between groups were performed using Student’s *t*-tests, and *p* values are presented. **C** The relative abundances of WT and variant *FOXL2* mRNA were analyzed in KGN and COV434 cells by pyrosequencing (left graph), allele-specific RT-PCR (middle graph) and real-time RT-PCR (right graph). gDNA was detected as a positive control. The relative abundances of the variant *FOXL2* mRNA were normalized to that of WT mRNA (set to 1). *FOXL2* mRNA levels detected by real-time RT-PCR were normalized to matching gDNA levels. The pyrosequencing data are presented from two independent experiments. The allele-specific semi-quantitative and real-time RT-PCR data are presented as the mean ± SEM from three independent experiments. \*\*\**p* < 0.001. n.d. not detected. **D** RNA-decay rates of WT and 402C>G *FOXL2* mRNAs in KGN cells were determined after treatment with 5 µg/mL ActD for the indicated times. The estimated half-lives of each transcript are presented. The data are presented as the mean ± SEM from three independent experiments.

We obtained analogous results using AGCT-derived KGN cells by pyrosequencing, allele-specific semi-quantitative RT-PCR, allele-specific real-time RT-PCR, and primer extension assays, which are heterozygous for the 402C>G mutation (Fig 1C, Appendix Fig S1G, and S1H). The relative abundance of variant *FOXL2* mRNA was ∼22% of WT *FOXL2* mRNA levels, while the gDNA levels of both alleles were similar in KGN cells (Fig 1C). When COV434, a cell line derived from JGCTs lacking the 402C>G mutation (Jamieson, Butzow *et al*., 2010), was tested as a control using the mutant-allele specific primer, no amplicons containing the 402C>G mutation were detected (Fig 1C and Appendix Fig S1G). We also observed a comparable allelic imbalance by allele-specific real-time RT-PCR on full-length *FOXL2* mRNA generated by the cap analysis of gene expression (CAGE) method from KGN cells (Appendix Fig S1I).

Next, we tested whether the decreased steady-state levels of the variant *FOXL2* mRNA resulted from alterations in mRNA stability. WT and variant *FOXL2* mRNA abundances were measured in KGN cells at several time points using allele-specific real-time RT-PCR after adding actinomycin D (ActD), which blocks transcription. The level of variant *FOXL2* mRNA decreased more rapidly than that of WT mRNA (t_1/2_ = 3.68 h for variant *FOXL2* versus t_1/2_ = 15.43 h for WT), indicating that the lower steady-state level of the variant *FOXL2* mRNA resulted from decreased mRNA stability (Fig 1D). Variant *FOXL2* mRNA instability was unlikely due to its altered secondary structure, as the 402C>G mutation was not predicted to affect the mRNA structure, based on M-fold analysis (http://mfold.rna.albany.edu/?q=mfold/RNA-Folding-Form) (Zuker, 2003) (Appendix Fig S2A and S2B).

### Selective degradation of the variant *FOXL2* mRNA by miR-1236

Next, we determined whether selective degradation of the variant *FOXL2* mRNA was due to miRNAs targeting the mutation site in the CDS. By analyzing the variant *FOXL2* mRNA sequence with an miRNA-prediction tool (genie.weizmann.ac.il/pubs/mir07/mir07_prediction.html) (Kertesz, Iovino *et al*., 2007), we identified miRNAs predicted to bind the sequence surrounding the mutation (Appendix Table 1). We selected five miRNAs predicted to preferentially bind the mutant allele over the WT allele for further analysis. Using DNA oligonucleotides complementary to these miRNAs (anti-miRNAs), which effectively inhibited the respective miRNAs based on upregulation of their known target mRNAs (Appendix Fig S3A), we found that anti-miR-1236 specifically increased the variant *FOXL2* mRNA-expression level without affecting that of WT *FOXL2* in KGN cells (Fig 2A). The remaining four anti-miRNAs did not affect the mRNA levels of WT *FOXL2* or the variant (Fig 2A). Consistent with this effect at the mRNA level, anti-miR-1236 also increased FOXL2 protein expression (Fig 2B). Conversely, transfection of an miR-1236 RNA mimic decreased FOXL2 protein expression (Fig 2C). In contrast to our observations with KGN cells, transfecting miR-1236 into COV434 cells (which lack the 402C>G mutation) did not affect FOXL2 protein expression (Fig 2C). To further validate the specificity of miR-1236 on the *FOXL2* variant, we cotransfected a WT- or variant FOXL2-expression plasmid together with miR-1236 RNA into 293T cells expressing minimal WT FOXL2 (but not the variant), and changes in FOXL2 expression were monitored by western blotting. The miR-1236 mimic specifically decreased the variant FOXL2 level without affecting that of WT FOXL2 (Appendix Fig S3B). The effective depletion or overexpression of miR-1236 in cells transfected with anti-miRNAs or mimic oligonucleotides, respectively, was confirmed with a TaqMan® microRNA assay for miR-1236 (Appendix Fig S3C).

**Figure 2.**
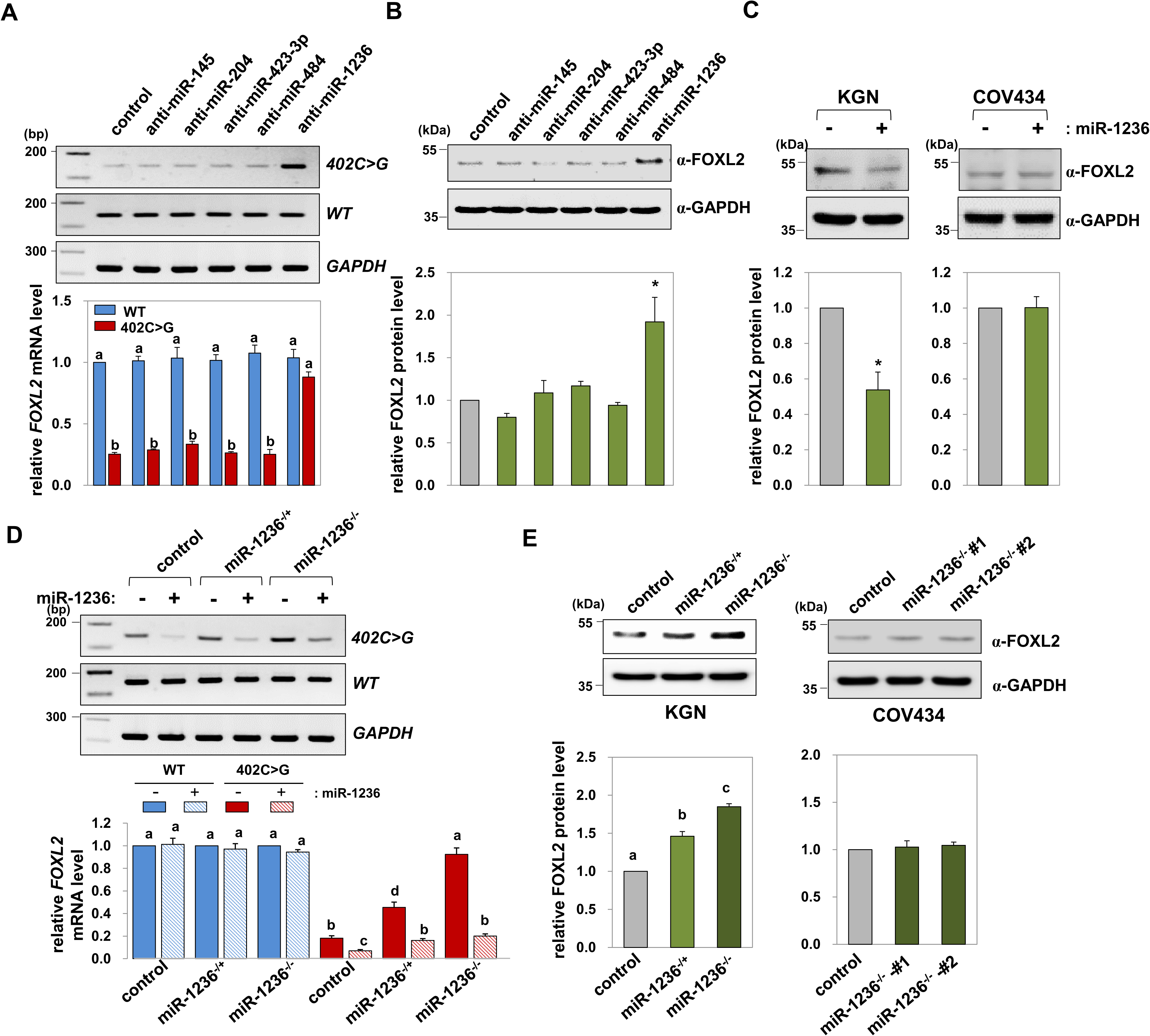
Selective downregulation of the variant *FOXL2* mRNA allele by miR-1236. **A, B** Changes in WT and variant *FOXL2* mRNA expression were assessed by RT-PCR (top) and real-time RT-PCR (bottom) (A) or by western blot analysis (B) after transfecting KGN cells with anti-miRNAs for 48 h. **C** FOXL2 protein levels after transfection of control or miR-1236 were assessed in KGN and COV434 cells. **D** The mRNA levels of WT and variant *FOXL2* in control, miR-1236^−/+^, and miR-1236^−/−^ KGN cells, with or without miR-1236 transfection, were determined by RT-PCR (top) or real-time RT-PCR (bottom). **E** FOXL2 protein expression in control, miR-1236^−/+^, and miR-1236^−/−^ KGN cells, and two independent miR-1236^−/−^ (#1 and #2) COV434 cell lines were determined by western blotting. Representative gel images are also shown. All quantified results (mean ± SEM) are from at least three independent experiments. Different letters (*p* < 0.0001) or asterisks (\**p* < 0.05) denote significant differences.

Moreover, we generated miR-1236-knockout (KO) KGN and COV434 cell lines using a clustered regularly interspaced short palindromic repeat (CRISPR)/CRISPR-associated protein 9 (Cas9)-nickase-based system (Appendix Fig S4A–4H) that is known to exert minimal off-target effects (Ran, Hsu *et al*., 2013). Using extracted total RNA from the KO cells, we performed TaqMan® microRNA analyses and confirmed miR-1236 depletion (Appendix Fig S4B and S4F). Northern blotting showed that miR-1236 was detectable in control KGN cells, but not in miR-1236^−/−^ cells (Appendix Fig S4I). Then, we evaluated WT and variant *FOXL2* mRNA levels in the KO cells. Variant *FOXL2* mRNA expression reverted to the level of WT mRNA when both alleles (miR-1236^−/−^) were excised and was partially restored when a single allele (miR-1236^−/+^) of miR-1236 was disrupted, without affecting WT *FOXL2* mRNA expression (Fig 2D). Re-introducing miR-1236 mimic via transfection in miR-1236-KO cells downregulated variant *FOXL2* mRNA expression without altering WT *FOXL2* expression (Fig 2D), demonstrating that restoration of variant *FOXL2* mRNA level resulted from miR-1236 KO. The observation that known miR-1236 target genes, *AFP* and *ZEB1*(Gao, Cai *et al*., 2015, Wang, Yan *et al*., 2014), were upregulated in these KO cells also suggested that efficient and effective excision of miR-1236 occurred (Appendix Fig S4J and S4K). Consistently, elevated FOXL2 protein expression was confirmed in miR-1236-KO KGN cells, whereas no change in FOXL2 expression occurred in miR-1236-KO COV434 cells (Fig 2E). These *in vivo* genome-editing results further corroborated miR-1236 as an endogenous functional miRNA that selectively acted on variant *FOXL2* mRNA.

### Allele-specific downregulation of the *FOXL2* mRNA variant with miR-1236 via targeting the 402C>G mutation site

Reporter constructs were developed to confirm that miR-1236 targets the 402C>G locus in the variant *FOXL2* mRNA CDS. A 231-bp DNA fragment containing the 402C>G locus of the *FOXL2* variant or the corresponding fragment of WT *FOXL2* was inserted, in-frame, into the CDS of the luciferase gene (Fig 3A). miR-1236 transfection decreased the activity of the luciferase reporter harboring the 402C>G *FOXL2* variant sequence by 70%, without affecting the activity of the luciferase reporter containing the WT *FOXL2* sequence (Fig 3B). Similarly, we observed allele-specific repression when the 231-bp *FOXL2* fragments were inserted in the 3′-UTR of the luciferase reporter gene (Fig 3C and 3D). The specificity of the interaction between miR-1236 and the variant locus in *FOXL2* mRNA was further tested using two miR-1236 mutants. The miR-1236-G mutant shifted the 7-mer seed match from the 402C>G *FOXL2* mRNA to the WT *FOXL2* mRNA (Fig 3E), whereas the seed sequence of the miR-1236-U mutant did not match either 402C>G or WT *FOXL2* mRNA (Fig 3F). Notably, the selectivity of the miR-1236-G mutant was reversed, as it repressed the WT reporter without affecting the 402C>G mutant reporter (Fig 3E). In contrast, the miR-1236-U mutant did not suppress the 402C>G mutant or WT *FOXL2* reporter (Fig 3F).

**Figure 3.**
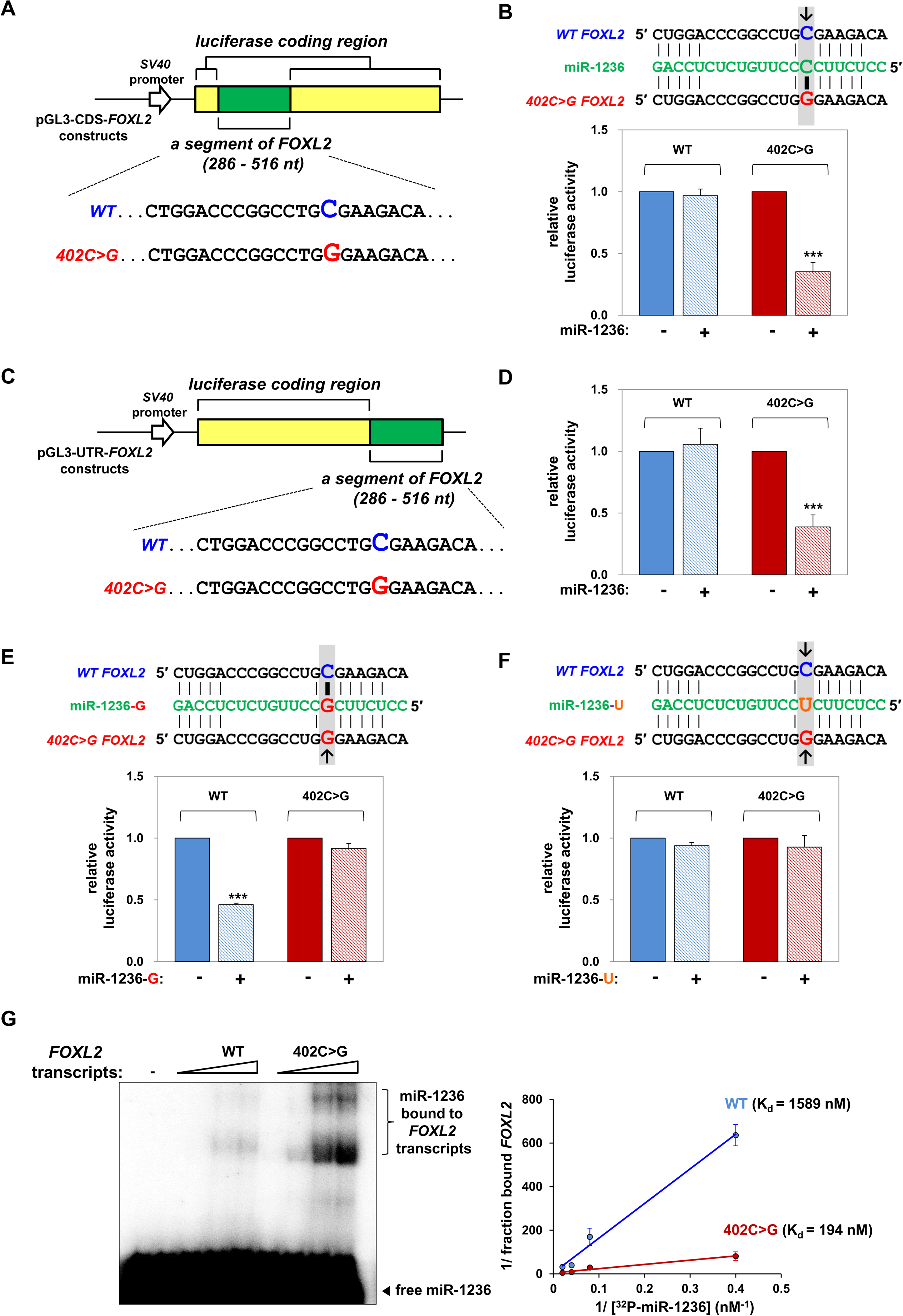
Allele-specific downregulation of *FOXL2* mRNA by *miR-1236*, which targets 402C>G. **A** Schematic representation of the luciferase-reporter constructs used to assay miR-1236 activity against a CDS target site in *FOXL2* mRNA. The 231-bp human *FOXL2* segments harboring either the C402 (WT) or the G402 (mutant) nucleotide were inserted in-frame into the CDS of the *luciferase* gene in the pGL3 control vector. **B** Luciferase activity of the reporter constructs shown in (A) were measured in KGN cells after transfection with an miR-1236 mimic for 48 h. **C** A schematic diagram of the luciferase-reporter constructs generated by inserting the predicted miR-1236-target sequences of WT and 402C>G *FOXL2* mRNAs in the 3′-UTR of *luciferase*. **D** Luciferase activities were measured in KGN cells, using the reporter constructs shown in (C), after transfection with a control miRNA or an miR-1236 mimic for 48 h. **E, F** miR-1236 mutants, in which the C that pairs with G402 of the *FOXL2* mutant was substituted with either G (miR-1236-G) (E) or U (miR-1236-U) (F), were cotransfected into KGN cells with one of the reporter constructs described above. Luciferase activities were subsequently determined. Arrows indicate the mismatched sites. **G** *In vitro* annealing kinetics of miR-1236 with 230 nt-long transcripts of WT or variant *FOXL2*. ^32^P-labeled miR-1236 (0.5 nM) was incubated with increasing concentrations of synthetic *FOXL2* transcripts (0, 2.5, 12.5, 25, or 50 nM). *FOXL2* mRNA–miR-1236 complexes were resolved on a 6% native gel and detected by autoradiography (left). The predicted K_d_s for the WT and 402C>G *FOXL2* transcripts are presented in the right graph. The data are expressed as the mean ± SEM from three independent experiments, performed in triplicate. \*\*\**p* < 0.001

In addition, the preferential binding of miR-1236 to 402C>G over the WT *FOXL2* transcript was confirmed by performing *in vitro-*binding assays. Synthetic 230-nt transcripts of WT or 402C>G *FOXL2* mRNA were incubated with radiolabeled miR-1236, and RNA-duplex formation was monitored. miR-1236 preferentially duplexed with the 402C>G *FOXL2* transcript, with predicted dissociation constants (K_d_) of 1,589 and 194 nM for the WT and 402C>G *FOXL2* transcripts, respectively (Fig 3G). Thus, these data indicate that the 402C>G locus was critical for distinguishing the effects of miR-1236 on *FOXL2* expression.

### AGO3 as the major miRISC component for miR-1236-mediated *FOXL2* variant mRNA degradation

Because AGO2 is known to act primarily on the 3′-UTR of target mRNAs (Hafner *et al*., 2010) and a recent study demonstrated that AGO3 is also associated with slicer activity (Park *et al*., 2017), we investigated the possibility that AGO3 can regulate RISC activity against the variant *FOXL2* mRNA by recognizing the mutated site in its CDS. Each AGO was knocked down using siRNAs, and changes in the levels of WT and variant *FOXL2* mRNAs were examined in KGN cells. Of interest, we found that AGO3 knockdown preferentially increased mRNA expression of the *FOXL2* variant without affecting that of WT *FOXL2* (Fig 4A). In contrast, AGO2 knockdown increased both the WT and variant mRNAs (Fig 4A), indicating that WT *FOXL2* mRNA is degraded by AGO2-mediated miRNAs targeting the 3′-UTR, as previously described (Dai, Sun *et al*., 2013, Luo, Wu *et al*., 2015, Wang, Li *et al*., 2015). Depletion of either AGO1 or AGO4 did not affect the levels of the WT and variant *FOXL2* mRNAs (Fig 4A). Consistent with the effects on the mRNA levels, increased FOXL2 protein levels were observed in KGN cells following the depletion of either AGO2 or AGO3 (Fig 4B). To determine whether miR-1236 mediated these effects, AGO-depleted KGN cells were transfected with miR-1236, and changes in *FOXL2* mRNA-expression levels were evaluated by allele-specific real-time RT-PCR. miR-1236 overexpression did not alter WT *FOXL2* mRNA expression in cells with AGO1, AGO2, AGO3, or AGO4 depletion (Fig 4C; left graph). In contrast, miR-1236 transfection downregulated marginally variant *FOXL2* mRNA expression in AGO3-knockdown cells (Fig 4C; right graph). miR-1236 transfection was partially effective in AGO2-knockdown cells, but efficiently downregulated variant *FOXL2* mRNA levels in AGO1- or 4-depleted cells (Fig 4C; right graph). In addition, to ascertain the role of AGO3 in the activity of miR-1236 in promoting variant *FOXL2* mRNA decay, changes in *FOXL2* mRNA-expression levels were determined in miR-1236-KO cells after silencing each AGO. Our results were similar to those shown in Fig 4A when the control cell line expressing miR-1236 (miR-1236^+/+^) was used in these experiments (Fig 4D; left graph). In sharp contrast, depletion of AGO1, 3, or 4 failed to increase the level of variant *FOXL2* mRNA in miR-1236-KO cells (miR-1236^−/−^) (Fig 4D; right graph). Consistent with the results shown in Fig 4A, AGO2 depletion increased both WT and variant *FOXL2* mRNA expression in miR-1236^−/−^ cells (Fig 4D; right graph).

**Figure 4.**
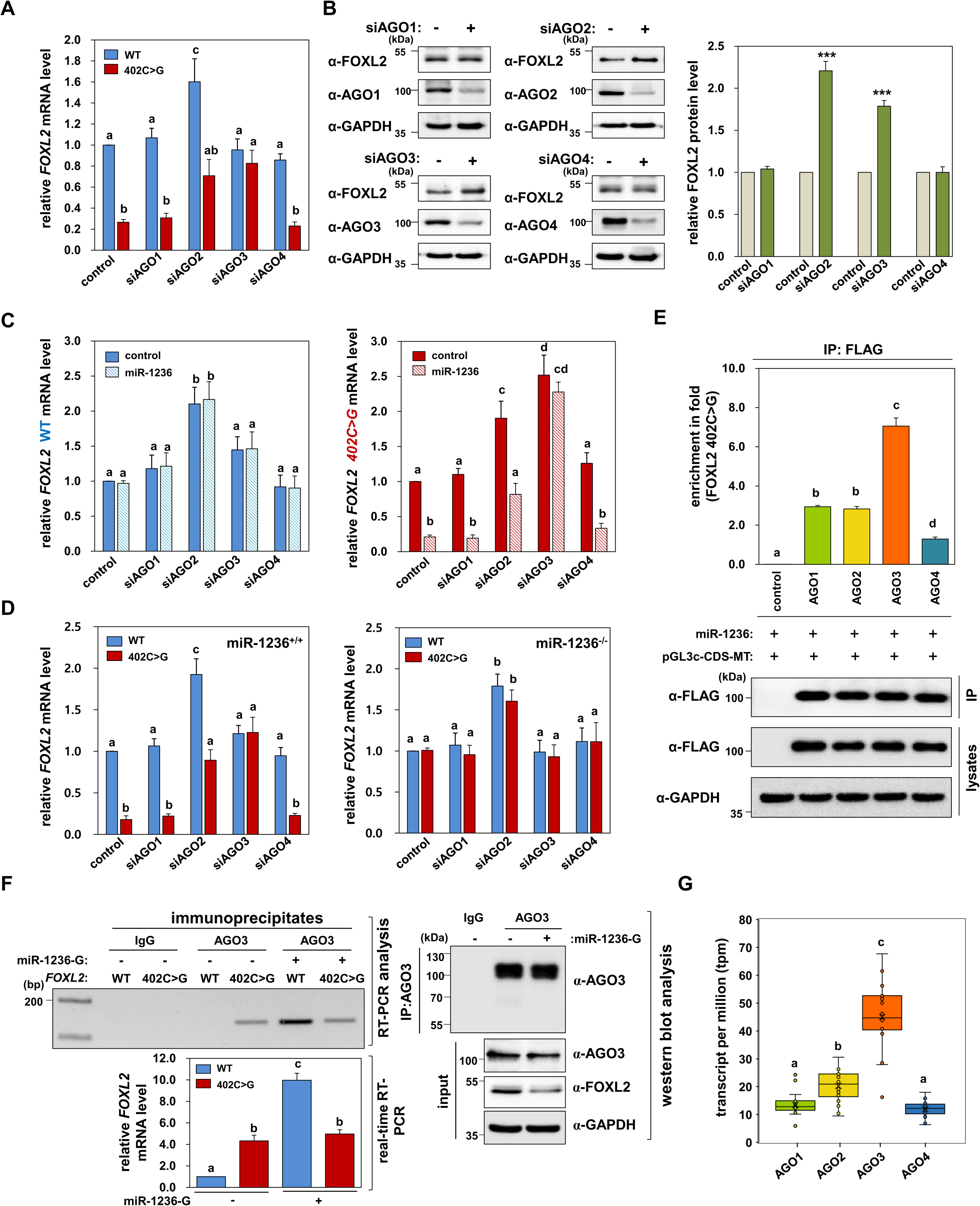
Identification of AGO3 as the major miRISC regulator for variant *FOXL2* mRNA degradation. **A**, **B** Changes in WT and variant *FOXL2* mRNA-expression levels were assessed by real-time RT-PCR (A) or western blot analysis (B) after transfecting KGN cells with siRNAs against AGO mRNAs for 48 h. The data (mean ± SEM) are from three independent experiments, performed in triplicate. **C** The mRNA levels of WT (left) and variant *FOXL2* (right) were determined in KGN cells by real-time RT-PCR, after transfecting a control miRNA or miR-1236. The data (mean ± SEM) are from three independent experiments, performed in triplicate. **D** The mRNA levels of WT and the variant *FOXL2* in control (left) and miR-1236*^−/−^* KGN cells (right) after transfecting siRNAs against AGO mRNAs were determined by allele-specific real-time RT-PCR. The data (mean ± SEM) are from three independent experiments, performed in triplicate. **E** 293T cells were transfected with an miR-1236 mimic (50 nM) for 24 h, followed by cotransfection for 24 h with expression vectors encoding FLAG/HA-tagged variants of the indicated human AGOs and pGL3c-*FOXL2*-CDS. The empty p3XFLAG-CMV-10 vector was used as control. Co-immunoprecipitated mRNAs were reverse transcribed, and the cDNA products were used for allele-specific real-time PCR analysis of the *FOXL2* variant (top). The immunoprecipitated-AGO proteins were detected by western blotting (bottom). The data (mean ± SEM) are from three independent experiments. **F** *In vivo* association of AGO3-mediated miRISC formation with *FOXL2* mRNAs are shown. Following transfection of a control miRNA or the miR-1236-G mutant into KGN cells, AGO3-mediated RISC-associated RNAs were isolated by immunoprecipitation with an anti-AGO3 antibody. IgG was used as a control. The co-immunoprecipitated mRNAs were reverse transcribed using a *FOXL2*-430-R primer binding downstream of the 402C>G site. The cDNA products were used for *FOXL2* allele-specific PCR analysis with the FOXL2-279F primer (Appendix Fig. S1), and a representative result (top left) is shown. Quantified real-time RT-PCR results (bottom left) are also presented. Western blot analysis of immunoprecipitated AGO3 and inputs are shown in the right panel. The data are presented as the mean ± SEM of two independent experiments. **G** RNA-seq analysis was performed to determine AGO-expression levels (transcripts per million) from the individual tissues from 20 independent AGCT patients. Different letters (*p* < 0.05) or asterisks (\*\*\**p* < 0.001) denote statistically significant differences.

To assure miRISC formation between AGO3 and miR-1236, the level of enriched miR-1236 in KGN cell immunoprecipitates with each AGO was determined following transfection of FLAG-tagged AGOs, miR-1236, and a pGL3c-CDS-variant *FOXL2* construct (Fig 3A). We found that variant *FOXL2* mRNA was highly enriched in the AGO3 immunoprecipitate compared to the other AGO immunoprecipitates (Fig 4E). We further detected the *in vivo* formation of an miRISC comprised of endogenously expressed variant *FOXL2* mRNA, miR-1236, and AGO3. The variant *FOXL2* transcript was highly incorporated into the AGO3 immunoprecipitate, whereas negligible WT *FOXL2* was incorporated (Fig 4F). Preferential incorporation of the variant *FOXL2* transcript with AGO3 was reversed by transfection of the miR-1236-G mutant into KGN cells, in which predominant incorporation of WT *FOXL2* mRNA occurred with a concomitant decrease in the FOXL2 protein level (Fig 4F). Together, these results indicate that a functional miRISC consisting of AGO3, miR-1236, and the variant *FOXL2* mRNA preferentially formed in KGN cells.

KGN cells express all four AGOs, and a comparison of their relative abundances indicated that *AGO3* was the most abundant (Appendix Fig S5). Further, we analyzed the relative abundances of *AGO* mRNAs in individual AGCT tissues from patients, using high-throughput deep sequencing. As shown in Fig 4G, we observed dramatically higher expression of *AGO3* mRNA-expression levels compared to those of *AGO1*, *AGO2*, and *AGO4* in 20 independent patients with AGCT, supporting a previously unidentified role of AGO3 as a functional miRISC component in AGCT cells.

### Identification of DHX9 as a functional component for AGO3–miRISC regulation of variant *FOXL2* mRNA

We further investigated potential functional AGO3–miRISC components that could regulate expression of the variant *FOXL2* mRNA. Since most miRNAs target in CDS regions in plants(Iwakawa & Tomari, 2013), candidate human proteins were selected based on searches for homologous genes in plants that regulate the activities of miRISCs, AGO2-binding proteins, and RNA-binding proteins. Among them, silencing DHX9, an ATP-dependent RNA helicase A that has been reported to function as a siRNA-loading and-recognition factor for AGO2-siRISC assembly (Fu & Yuan, 2013, Robb & Rana, 2007), prominently affected the abundance of variant *FOXL2* mRNA (Fig 5A). In particular, DHX9 silencing restored the variant *FOXL2* mRNA level to that of the WT mRNA, involving a 6-fold increase of variant *FOXL2* mRNA compared to a 1.7-fold increase of the WT mRNA (Fig 5A). In contrast, depletion of GW182, a well-known miRISC component that associates with AGO2, increased the abundances of both the WT and variant *FOXL2* mRNAs (Fig 5A). Consistent with these effects on the mRNA level, DHX9- or GW182-depletion also increased expression of the FOXL2 protein (Fig 5B). Immunoprecipitation analysis of AGOs with DHX9 or GW182 showed that DHX9 exhibited stronger binding affinity to AGO3, whereas GW182 bound AGO2 more tightly (Fig 5C). In addition, we determined the functional role of DHX9 for *in vivo* AGO3–miRISC formation with the variant *FOXL2* mRNA. Consistent with the data shown in Fig 4F, a predominant incorporation of the variant *FOXL2* transcript over the WT transcript in AGO3 immunoprecipitates was observed (Fig 5D). However, DHX9 depletion significantly decreased the incorporation of variant *FOXL2* mRNA in AGO3 immunoprecipitates without affecting the incorporation of WT *FOXL2* mRNA (Fig 5D). These results imply that DHX9 served as a critical factor required for AGO3–miRISC-associated miR-1236 function.

**Figure 5.**
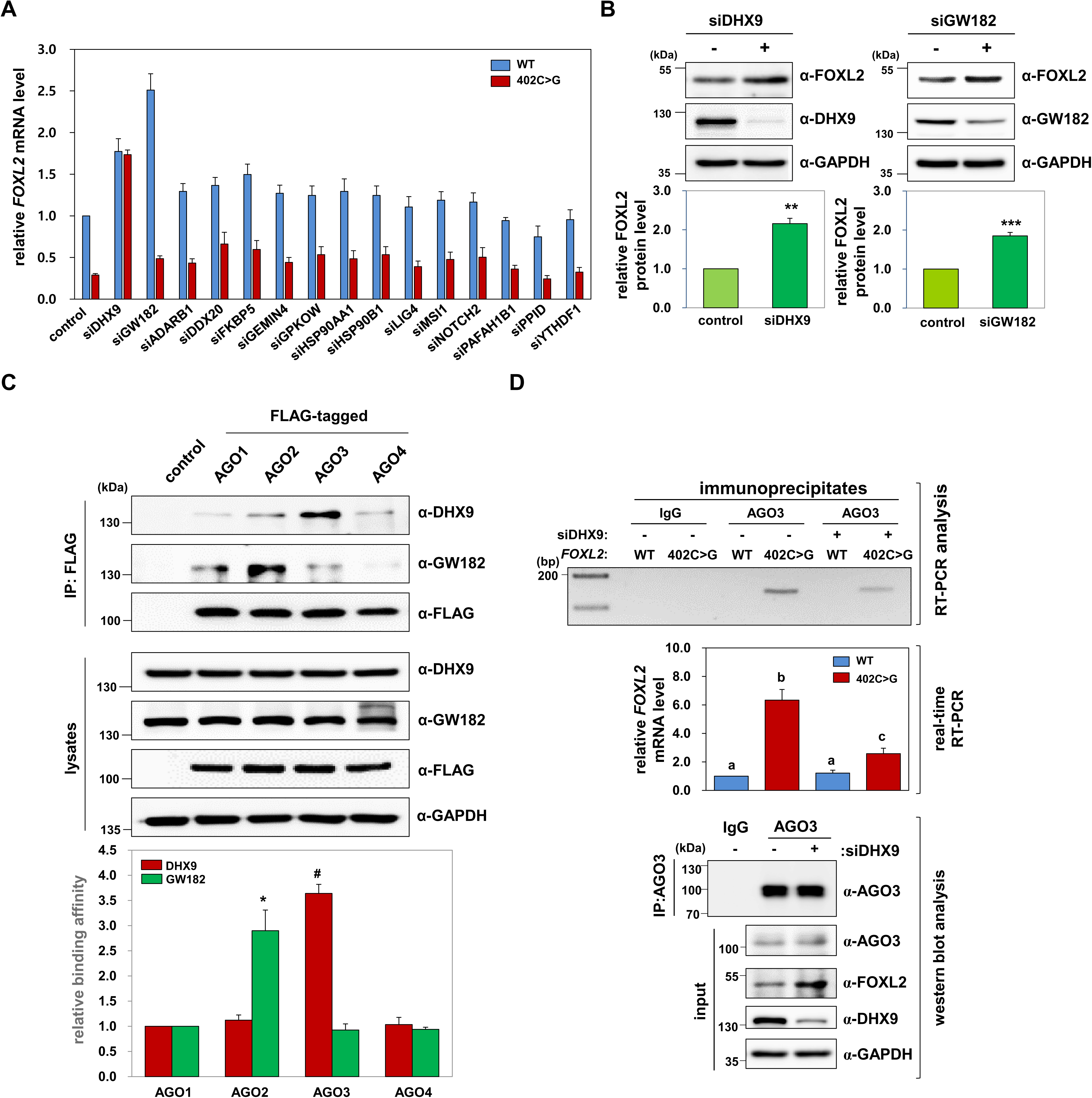
DHX9 as a preferential interactor with AGO3 and an essential molecule for AGO3-mediated degradation of the variant *FOXL2* mRNA. **A** Changes in WT and variant *FOXL2* mRNA levels in KGN cells were assessed by real-time RT-PCR after transfecting siRNAs against the indicated factors for 48 h. The data are presented as the mean ± SEM from three independent experiments, performed in triplicate. **B** FOXL2 protein-expression levels were determined by western blotting after transfecting KGN cells with control, DHX9, or GW182 siRNAs for 48 h. Quantification of FOXL2 protein expression is presented in the bottom panel. The data are presented as the mean ± SEM from three independent experiments. \*\**p* < 0.01, \*\*\**p* < 0.001. **C** Relative binding affinities of DHX9 and GW182 to AGOs. 293T cells were transfected with expression vectors encoding the indicated FLAG/HA-tagged AGOs, and cell extracts were prepared and immunoprecipitated with an anti-FLAG antibody, followed by immunoblot analyses. The empty p3XFLAG-CMV-10 vector was used as a control. The band intensities of immunoprecipitated DHX9 and GW182 were quantified and normalized following pull-down with the indicated AGOs. The data are presented as the mean ± SEM from three independent experiments. * and # indicate statistically significant differences in the respective amounts of DHX9 or GW182 bound to AGO1 (*p* < 0.05). **D** Following transfection of control siRNA or siDHX9 into KGN cells, AGO3-mediated RISC-associated RNAs were immunoprecipitated using an anti-AGO3 antibody. IgG was used as a control. Co-immunoprecipitated mRNAs were reverse transcribed using a *FOXL2*-430-R primer binding downstream of the 402C>G site. The cDNA products were used for *FOXL2* allele-specific PCR analysis with a FOXL2-279F primer (Appendix Fig S1A), and a representative result obtained by RT-PCR (top) is shown. Quantified real-time RT-PCR results (middle) are also presented. Western blots of immunoprecipitated AGO3 and the inputs are shown in the bottom panel. The data are presented as the mean ± SEM from three independent experiments.

### Oncogenic function of miR-1236 in AGCT cells

Next, we investigated the oncogenic effects of miR-1236 in AGCT cells. We found that transfecting KGN cells with an miR-1236 mimic or anti-miR-1236 significantly decreased or increased the numbers of annexin-V-positive apoptotic cells, and increased or decreased the cellular viability, respectively (Fig 6A, 6B, Appendix Fig S6A, and S6B). Overexpression of other anti-miRNAs, which showed no significant effects on *FOXL2* expression (Fig 2A and 2B), did not alter KGN cell viability (Appendix Fig S6C). Moreover, treatment with the miR-1236 mimic accelerated cell cycle progression to S phase, whereas anti-miR-1236 induced cell cycle arrest at G_0_/G_1_ phase (Fig 6C and 6D). Transfection of either miR-1236 or anti-miR-1236 in *FOXL2*-silenced cells failed to alter the numbers of apoptotic cells, cell cycle, and cell viability (Fig 6A-6D, Appendix Fig S6D, and S6E), indicating that the effects of miR-1236 on KGN cells were *FOXL2*-dependent. The effect of *FOXL2* on GCT cell migration was also examined using Transwell chambers, and we found that miR-1236 or anti-miR-1236 transfection promoted or suppressed cell migration, respectively, and these effects were abolished upon *FOXL2* depletion (Fig 6E and 6F). Transfecting miR-1236 into COV434 cells did not alter either cell viability or cell migration and had no effect on FOXL2 protein expression (Fig 6G and 6H), validating the 402C>G allele-specific oncogenic effects of miR-1236. Additionally, we confirmed the oncogenic properties of miR-1236 using miR-1236-KO KGN cells and obtained results that were consistent with the effects observed after anti-miR-1236 transfection. miR-1236^−/−^ KGN cells exhibited significantly reduced cell survival, proliferation, and migration compared to control KGN cells, and miR-1236^−/+^ KGN cells showed intermediate activities (Fig 6I, 6J, and 6K). In sharp contrast, the oncogenic characteristics were not altered in miR-1236*^−/−^* COV434 cells (Fig 6L).

**Figure 6.**
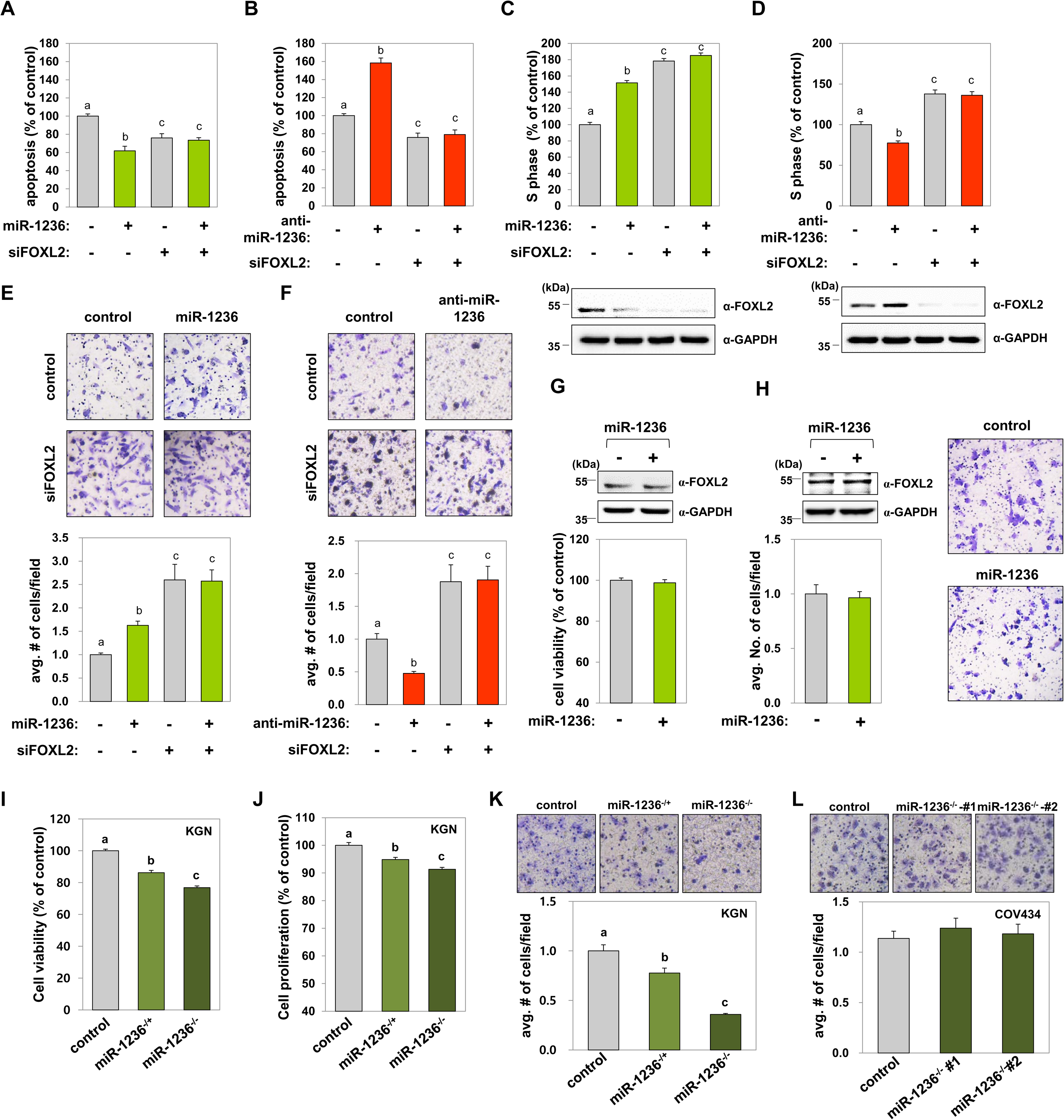
FOXL2-mediated oncogenic effects of miR-1236 on cell death, cell cycle progression, and cell migration. **A-D** KGN cells were transfected with 200 nM of scrambled control or FOXL2-specific siRNAs for 24 h. Then, KGN cells were further transfected with 20 nM of control miRNA, miR-1236, control anti-miRNA, or anti-miR-1236. The proportion of annexin-V-positive apoptotic cells (A and B) and the population at S phase (C and D) were analyzed by flow cytometry. The data are presented as the mean ± SEM of three independent experiments. Different letters denote statistically significant differences (*p* < 0.0001). Efficient silencing of *FOXL2* using specific siRNAs was confirmed by western blotting. The data are presented in the bottom panels as the mean ± SEM of three independent experiments, performed in triplicate. **E, F** KGN cells were transfected with control siRNA or si*FOXL2* for 24 h. Then, KGN cells were further transfected with control miRNA or miR-1236 (E) or with control anti-miRNA or anti-miR-1236 (F) for 48 h, and Transwell-migration assays were performed. The migrated cells were imaged under a bright-field microscope (100× magnification). The results are from three independent experiments and represent fold-changes in the average number of cells/field (mean ± SEM). Different letters denote statistically significant differences (*p* < 0.01). **G, H** COV434 cells were transfected with miR-control or miR-1236 for 48 h, after which cell viabilities (G) and migration abilities (H) were measured. Immunoblots showing no change in FOXL2 protein are presented in the top panel, and images of migrated cells are presented in the right panel. **I**-**K** The properties of miR-1236^−/+^ and miR-1236^−/−^ KGN cells versus control KGN cells were assessed by measuring cell viability (I), cell proliferation (J), and cell migration (K). **L** No difference in the cell-migration activities of control and two independent miR-1236^−/−^ (#1 and #2) COV434 cell lines. The migrated cells were imaged under a bright-field microscope (100× magnification; top) and the results (bottom) represent fold-changes in the average number of cells/field. The data are presented as the mean ± SEM of three independent experiments. Different letters denote statistically significant differences (*p* < 0.0001).

### *In vivo* suppression of metastasis by miR-1236 KO and inverse expression between miR-1236 and the variant *FOXL2* mRNA in tissues from AGCT patients

We further performed an *in vivo* xenograft mouse experiment using the miR-1236^−/−^ KGN cells and examined the effect of miR-1236 loss on AGCT metastasis. Xenografting control KGN cells resulted in AGCT metastasis to the intestine in mice (Fig 7A and 7B). In contrast, miR-1236^−/−^ cell-xenografted mice showed significantly fewer metastasized tumor nodules (Fig 7A and 7B). By analyzing the relative abundances of *FOXL2* transcripts in these tumors, we confirmed that the miR-1236-KO tumors expressed more variant *FOXL2* mRNA than WT mRNA (Fig 7C). These *in vivo* results provided further support that miR-1236 acts as a critical molecule for AGCT metastasis by preferentially targeting the variant *FOXL2* mRNA.

**Figure 7.**
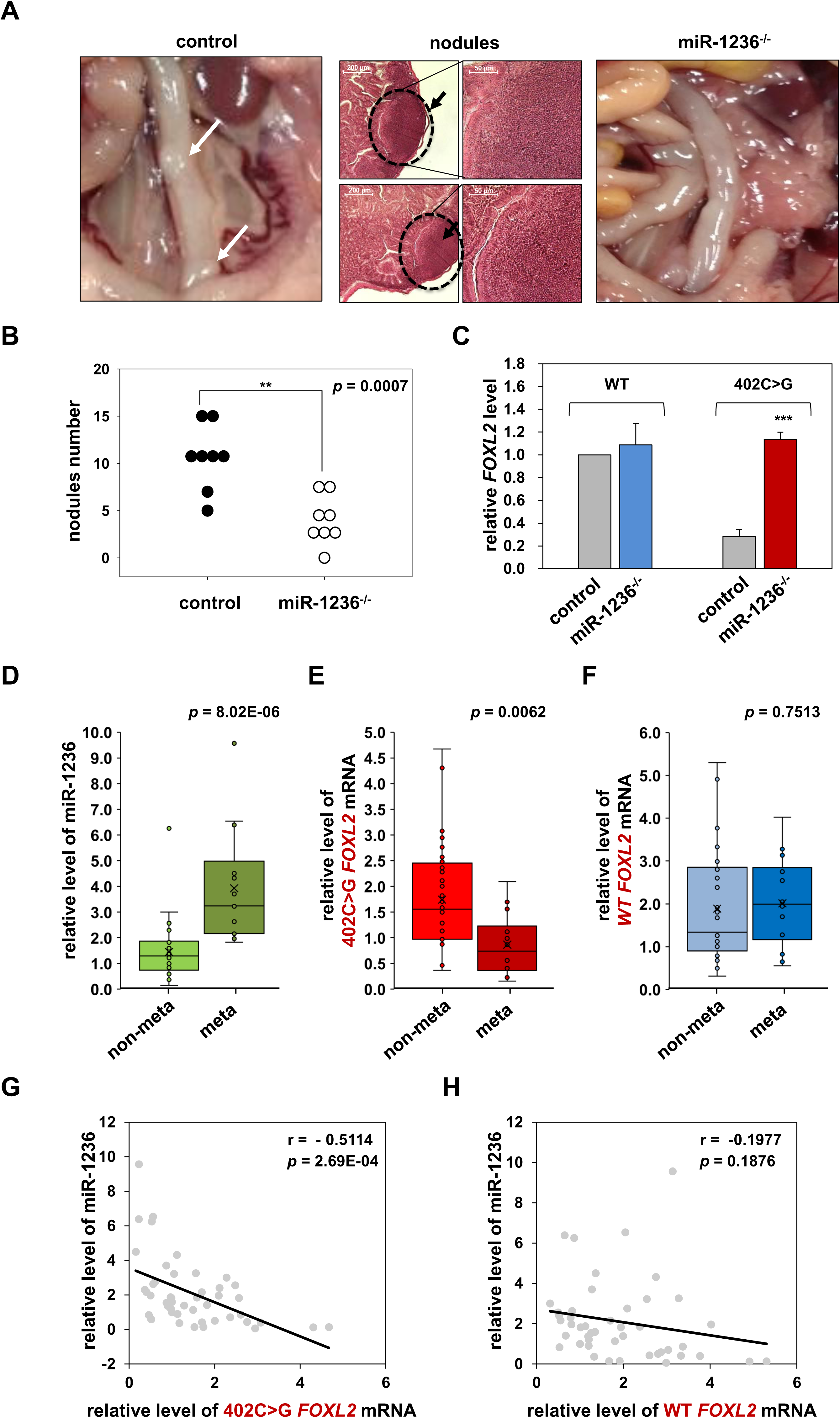
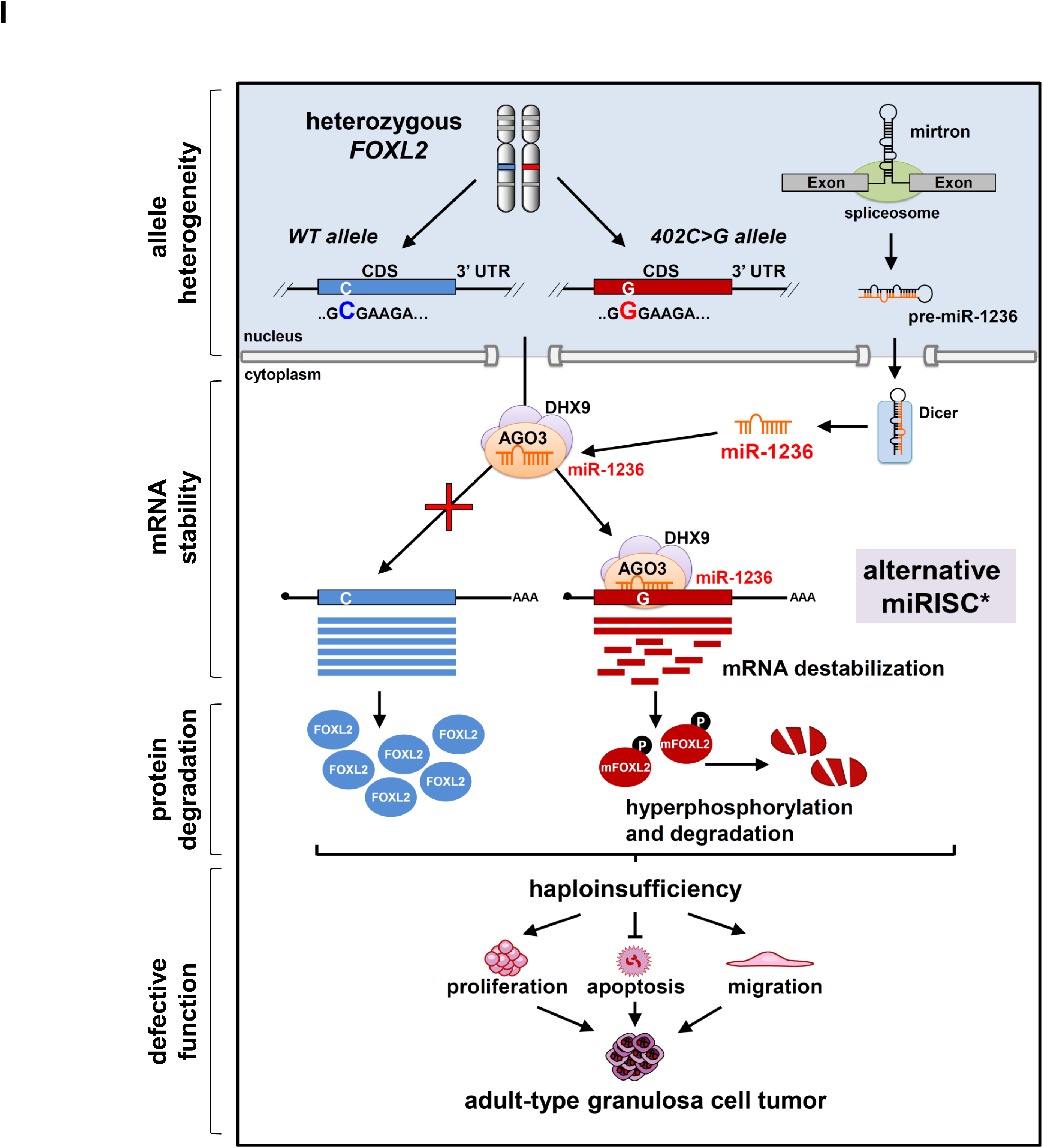
Inhibition of *in vivo* metastasis by miR-1236 KO and inverse expression between miR-1236 and the variant *FOXL2* mRNA in tissues from AGCT patients. **A**-**C** The effect of miR-1236 KO on AGCT metastasis was assessed, using an *in vivo* xenograft mice model. (A) Representative images of tumor nodules (white arrows) formed in the intestines of nude mice xenografted with control or miR-1236*^−/−^* KGN cells are shown (left). Hematoxylin and eosin staining confirmed the pathological characteristics of the metastasized GCT nodules (right, black arrows). (B) The number of tumor nodules formed in the intestines was counted in control (n = 8) and miR-1236^−/−^ (n = 8) mice. (C) Allele-specific real-time RT-PCR analysis of the WT and 402C>G variant *FOXL2* mRNAs was performed using RNA extracted from tumor nodules from control or miR-1236^−/−^ mice. \*\**p* < 0.01, \*\*\**p* < 0.001. **D-F** Box-and-whisker plots showing the relative expression of miR-1236 (D), variant *FOXL2* mRNA (E), and WT *FOXL2* mRNA (F), respectively, in 32 patients with non-metastasized AGCTs and 14 patients with metastasized AGCTs. The relative miR-1236 levels were measured using a TaqMan® microRNA RT-qPCR assay, with expression normalized to *RNU6B*. The levels of 402C>G or WT *FOXL2* mRNA were determined by real-time RT-PCR, and the data were normalized to paired-gDNA levels. The relative levels of miR-1236 and *FOXL2* mRNAs were quantified by setting the levels of AGCT #1 to 1. Real-time RT-PCR was performed in triplicate for each specimen. The box plot represents the lower, median, and upper quartiles, and the whiskers represent the 95% confidence interval of the mean. The whiskers extend to the most extreme data points not considered outliers, and the outliers are represented as dots. Comparisons between groups were performed using Student’s *t*-tests, and *p* values are presented. **G, H** The estimated regression line superimposed on the scatter plot of miR-1236 levels with 402C>G *FOXL2* mRNA (G) or WT *FOXL2* mRNA (H) in AGCT samples (n = 46) is shown, along with correlation coefficient (r) and *p* values. **I** The proposed model for *FOXL2* haploinsufficiency induced by the 402C>G mutation during AGCT development.

Moreover, we examined whether the aggressiveness of AGCT was indeed linked to the ability of miR-1236 to downregulate mRNA expression of the 402C>G *FOXL2* variant in patients with AGCT. Clinical tissues from AGCT patients (exhibiting metastasis or no evidence of metastasis) were examined for miR-1236 and *FOXL2* mRNA expression. Notably, increased miR-1236 and decreased 402C>G *FOXL2* mRNA levels were evident in metastatic AGCT tissues compared to non-metastatic AGCT tissues (Fig 7D and 7E). In contrast, the expression of WT *FOXL2* mRNA was similar in both AGCT groups (Fig 7F). Moreover, comparative analysis of the miR-1236 and *FOXL2* mRNA-expression levels in tissues from AGCT patients revealed a strong inverse correlation between miR-1236 and variant *FOXL2* mRNA expression (Fig 7G). Such a correlation was not observed between miR-1236 and WT *FOXL2* mRNA (Fig 7H). Together, these results provided further support that miR-1236 caused haploinsufficiency of the tumor-suppressor FOXL2 by inducing degradation of the variant *FOXL2* mRNA following recognition of the 402C>G locus as a target site, suggesting that miR-1236 plays an important etiological role in AGCT (Fig 7I).

## DISCUSSION

Data from recent studies have suggested that miRNAs can target CDS sites in human cells, and the presence of functional miRNA recognition elements in CDSs was addressed in a recent human study (Cai, Cao *et al*., 2015, Duursma, Kedde *et al*., 2008, Elcheva *et al*., 2009, Forman *et al*., 2008, Schnall-Levin *et al*., 2011, Wang, Long *et al*., 2012, Zhang *et al*., 2018). However, the regulatory mechanism underlying miRNA activity against the CDS of target genes is largely unknown. The general conceptual agreement that miRNAs target 3′-UTRs mostly arises from the idea that translating ribosomes occupy the CDS region and interfere with RISC formation. Thus, the binding of a miRISC to the CDS region of a transcript has been considered a less efficient mechanism of action than binding to the 3′-UTR (Bartel, 2009, Gu, Jin *et al*., 2009). However, a recent report showed that RISC formation in the CDS region causes translation repression, but do not appear to alter the overall ribosome occupancy on the target mRNA (Zhang *et al*., 2018), indicating the existence of disparate mechanism for translation repression of target mRNA by CDS-targeted miRNAs. In addition, over 70% of mammalian genes generate mRNA transcripts with differing lengths and 3′-UTR sequences due to alternative cleavage and polyadenylation, which would abolish some miRNA-binding sites (Tian & Manley, 2017). Accordingly, variations in 3′-UTR sequences would decrease the efficiency of miRNA-mediated target mRNA degradation, which is often observed with oncogenes in cancer (Mayr & Bartel, 2009, Mayr, Hemann *et al*., 2007). Therefore, targeting CDSs can assure effective downregulation of gene expression by miRNA, especially for disease-associated genes whose expression levels need to be tightly controlled.

miR-1236, the trans-acting RNA regulator identified in this study, is a mirtron located in an intron of the negative-elongation factor complex, member E (*NELFE*) gene (Okamura, Hagen *et al*., 2007, Ruby, Jan *et al*., 2007, Wen, Ladewig *et al*., 2015). The genomic sequences of the miR-1236 region are well conserved from lower vertebrates (including cavefish, lizards, and chickens) to mammalian vertebrates (Appendix Fig S7). The steady state level of miR-1236 expression seems to be more than 300 copies per KGN cell, based on northern blot analysis of small RNAs purified from 2 × 10^7^ *miR-1236^+/+^* KGN cells, which showed a radioactive intensity similar to 10 fmol of miR-1236 mimics (Appendix Fig S4I). According to a very recent miRNA profiling data by Kim *et al* (Kim, Kim *et al*., 2019), in which they developed and adopted bias-minimized accurate quantification by sequencing (AQ-seq) technology, miR-1236 was fairly well expressed as being in the top 50% of abundance among all miRNAs detected in human cells. Cumulative recent data revealed altered various cellular responses after modulating miR-1236 expression levels in diverse cell types (An, Ma *et al*., 2019, Chen, Teng *et al*., 2016, Gao *et al*., 2015, Jones, Li *et al*., 2012, Ma, Shen *et al*., 2014, Sato, Yoshimura *et al*., 2012, Thoma, 2015, Wang, Tang *et al*., 2016, Wang *et al*., 2014, Wang, Liu *et al*., 2018, Zhu, Wang *et al*., 2018). In this study, miR-1236 selectively targeted the mutated 402C>G locus in the *FOXL2* CDS, without affecting WT *FOXL2* mRNA expression. Preferential targeting of the mutated ‘G’ allele by miR-1236 was also reported in recent single-nucleotide polymorphism (SNP) studies: specific miR-1236 binding to the G allele over the C allele (rs11536889) in the 3′-UTR of Toll-like receptor 4 (Zhao, Feng *et al*., 2019) and specific miR-1236 targeting of the G allele over the T allele (rs4246215) in the 3′-UTR of Flap endonuclease 1 (Nanda, Kumar *et al*., 2018). Furthermore, our *in vivo* and *in vitro* analysis of the phenotypes of miR-1236 KO cells collectively showed that miR-1236 acted as an oncomiR in AGCT cells by causing selective variant *FOXL2* mRNA degradation (Fig 6 and 7).

For miR-1236 activity against variant *FOXL2* mRNA, we identified AGO3 as a key AGO protein (Fig 5). For over two decades, AGO2 has been considered as the key AGO protein in miRISC formation. However, AGO3 may have a distinct, but overlapping functionality for mRNA degradation, considering its slicer ability, the presence of the same conserved motif in the catalytic center for mRNA cleavage, and its ability to interact with a similar spectrum of RNA transcripts (Azuma-Mukai *et al*., 2008, Huang, Li *et al*., 2014, Landthaler, Gaidatzis *et al*., 2008, Meister *et al*., 2004, Park *et al*., 2017). This study provides evidence supporting this notion, considering that AGO3 depletion completely restored the variant *FOXL2* mRNA-expression level to the WT level; that an *in vivo* miRISC was comprised of endogenously expressed variant *FOXL2* mRNA, miR-1236, and AGO3; and that the abundance of AGO3 was highest among the AGOs in AGCT tissues (Fig 4). Thus, we conjecture that the association of AGO3–RISC with target RNAs could be stabilized or promoted by interactions with distinct binding partner proteins, thereby facilitating target mRNA decay in the CDS. We discovered DHX9 as one of such proteins, which interacted preferentially with AGO3 (Fig 5C). DHX9 depletion increased variant *FOXL2* mRNA expression to the WT level, and variant *FOXL2* association with AGO3 decreased markedly in DHX9-depleted cells (Fig 5A, 5B, and 5D). Compromised target mRNA incorporation and processing was not due to inefficient miR-1236 incorporation into AGO3–miRISC after DHX9 depletion (Appendix Fig S8). In contrast to effects of DHX9 depletion, depletion of GW182, of which association with AGO2 is essential for gene silencing by miRNAs in animals (Eulalio, Tritschler *et al*., 2009), increased the abundances of both the WT and variant *FOXL2* mRNAs (Fig 5A and B), indicating that GW182 is required for miRISC action on the 3′-UTR present in both *FOXL2* mRNAs. In addition, GW182 preferentially bound AGO2 (Fig 5C), suggesting the presence of preferential binding partners for each AGO. While the detailed regulatory mechanism by which this non-canonical miRISC involving AGO3 and DHX9 targets the variant *FOXL2* CDS needs to be studied further, it is worthwhile to note findings of a recent study showing that CDS-targeted miRNAs require extensive base-pairing at the 3′ side rather the 5′ seed in a manner independent of GW182 expression (Zhang *et al*., 2018), which overlap with characteristics of miR1236.

Previously, we showed that the 134C>W missense mutant protein of FOXL2 caused by the 402C>G mutation in *FOXL2* induced serine 33 hyperphosphorylation by GSK3β, thereby accelerating ubiquitin-mediated proteasomal degradation of the mutant protein (Kim *et al*., 2014). Here, we demonstrated that the mutation dramatically destabilized variant *FOXL2* mRNA via the allele-specific activity of miR-1236. Our findings suggest an etiological model for *FOXL2*-mediated AGCT development; selective degradation of the variant *FOXL2* mRNA by an miRISC composed of AGO3, DHX9, and miR-1236 promotes a strong allelic imbalance of *FOXL2* expression, which is aggravated by accelerated degradation of the mutant FOXL2 protein. These processes result in FOXL2 haploinsufficiency and insufficient suppression of AGCT development (Fig 7I). In 2009, Shah *et al* reported RNA sequencing results of four AGCTs, in which they observed various ratios of the variant and WT *FOXL2* mRNA levels (9:1; 1:1, 4:6 and 6:4) (Shah *et al*., 2009). In the present study, we observed a dramatic decrease in the variant *FOXL2* mRNA levels compared to WT mRNA levels in 46 AGCT patients. Further studies are need to investigate how this discrepancy arises from. One possibility is that the RNA-seq study by Shah *et al* focused on expression of both *FOXL2* mRNAs rather than measurement of relative abundance of WT versus variant mRNAs. In fact, total sequencing and mapped reads in their study were not enough to accurately quantify gene expression. To obtain coverage over the full-sequence diversity of complex transcript libraries, including rare and low-expressed transcripts, up to 500 million reads are required (Fu, Xu *et al*., 2014). Our RNA-seq study generated, on average, 134,312,269 total sequence reads and approximately 71,531 Mbp of total mapped base using Illumina HiSeq 2500 for 20 AGCT samples. In addition, we employed three other methods to assess the relative abundance of the WT versus variant *FOXL2* mRNA and obtained a consistent allelic imbalance.

In conclusion, our study provides a pathogenic mechanism in which a disease-associated heterozygous missense mutation can induce strong degradation of a variant mRNA by an alternative miRISC, which largely contributes to the haploinsufficiency of the gene and, consequently, development of the associated disease. We identified miR-1236 as an oncogenic miRNA in AGCT, suggesting a promising therapeutic target and prognostic parameter for patients with AGCT. Our study also provides evidence of the importance of nucleotide substitutions, such as somatic mutations and SNPs, in the CDS in regulating gene expression at the level of mRNA stability. A research approach that diverges from focusing mainly on the effects of missense mutations on protein activity may uncover many previously unsuspected mechanisms involving the regulation of mRNA stability by phenotype-associated missense mutations. In addition, we identified miR-1236 as an oncogenic miRNA in AGCT, suggesting a promising therapeutic target and prognostic parameter for patients with AGCT.

## MATERIALS AND METHODS

### Cells culture and reagents

Human AGCT-derived KGN cells (Riken, Tsukuba, Japan) and JGCT-derived COV434 cells (Sigma-Aldrich, St. Louis, MO, USA) were cultured in Dulbecco’s modified Eagle’s medium (DMEM)-Ham’s F12 (Caisson, North Logan, UT, USA) and DMEM (Caisson), respectively. 293T cells (American Type Culture Collection) were cultured in DMEM. The media were supplemented with 10% fetal bovine serum (FBS) and 1% penicillin-streptomycin (Caisson). The cells were grown in 5% CO_2_ at 37°C. Other reagents were purchased from Sigma-Aldrich, unless otherwise indicated.

### Plasmids

Plasmids expressing Myc-tagged variants of WT FOXL2 and the C134W variant were prepared as described previously(Kim & Bae, 2014, Kim *et al*., 2011, Park, Shin *et al*., 2010). A 231-bp coding region (including nucleotide residue 402) of *FOXL2* containing the putative miRNA-recognition site was amplified from gDNA of KGN cells using the following primers (Cosmo Genetech, Seoul, Korea): CDS-F (5′-CATGCCATGGCAAAGGGCTGGCAAAATAG-3′), CDS-R (5′-CATGCCATGGGGCGCCTCCGGCCCCGAAGAG-3′), 3′-UTR-F (5′-GCTCTAGAAGGGCTGGCAAAATAGCA-3′), and 3′-UTR-R (5′-GCTCTAGACGGCGCCTCCGGCCCCGA-3′). The amplified fragment was then cloned into the pGL3-control vector (Promega, Madison, WI, USA) at the *Nco*I or *Xba*I sites (restriction enzymes from Takara Bio, Shiga, Japan) in the CDS or 3′-UTR of the luciferase gene, respectively, generating pGL3c-CDS-FOXL2s and pGL3c-UTR-FOXL2s. The mammalian expression vectors, FLAG/HA-tagged AGO1 (Addgene plasmid #10820), AGO2 (Addgene plasmid #10822), AGO3 (Addgene plasmid #10823), and AGO4 (Addgene plasmid #10824) were gifts from Dr. Thomas Tuschl(Meister *et al*., 2004).

### Human subjects and GCT tissues

Formalin-fixed paraffin-embedded (FFPE) block sections of AGCT (n = 46) specimens from patients who visited the Seoul Asan Medical Center and the Bundang CHA Women’s Hospital were analyzed. The present study was reviewed and approved by the Seoul Asan Medical Center and the Bundang CHA Women’s Hospital Institutional Review Board. Informed consent was obtained from all subjects who participated in this study. This study was carried out in compliance with approved guidelines.

### gDNA and RNA extraction

Total RNA from FFPE sections was extracted using the PureLink^TM^ FFPE Total RNA Isolation kit (Invitrogen, Carlsbad, CA, USA), following the manufacturer’s protocol. Total RNA in cultured cells was isolated using the TRIzol Reagent (Invitrogen), according to the manufacturer’s instructions. gDNA was extracted from the deparaffinized tissues and cells using an Intron G-DEX^TM^ Genomic DNA Extraction Kit (Intron, Seongnam, Korea), according to the manufacturer’s protocol.

### RNA library construction and ultra-deep RNA sequencing

Total RNA concentrations were calculated using Quant-IT RiboGreen (Invitrogen). To determine the % of RNA fragments >200 bp in length, samples are run on the TapeStation RNA ScreenTape (Agilent Technologies, Waldbronn, Germany). A sequencing library was constructed using 100 ng of total RNA and a TruSeq RNA Access Library Prep Kit (Illumina, Inc., San Diego, CA, USA), according to the manufacturer’s protocols. Briefly, total RNA was first fragmented into small pieces using divalent cations, at an elevated temperature. The cleaved RNA fragments were copied into first strand cDNA using SuperScript II reverse transcriptase (Invitrogen, cat#18064014) and random primers. Second-strand cDNA synthesis was performed using DNA Polymerase I, RNase H, and dUTP. The cDNA fragments were subjected to an end-repair process, involving the addition of a single ‘A’ base and then adapter ligation. The products were then purified and enriched by 15 cycles of PCR to create the cDNA library. All libraries were normalized, and six libraries were pooled into a single hybridization/capture reaction. The pooled libraries were incubated with a cocktail of biotinylated oligonucleotides corresponding to coding regions of the genome. Targeted library molecules were captured via hybridized biotinylated oligonucleotide probes using streptavidin-conjugated beads. After two rounds of hybridization/capture reactions, the enriched library molecules were subjected to a second round of PCR amplification in 10 cycles. The captured libraries were quantified using KAPA Library Quantification Kits for Illumina Sequencing platforms according to the qPCR Quantification Protocol Guide (Kapa Biosystems, catalog number KK4854) and qualified using the TapeStation D1000 ScreenTape assay (Agilent Technologies, catalog number 5067-5582). The indexed libraries were then loaded into an Illumina HiSeq 2500 instrument (Illumina, Inc.), and paired-end (2 × 100 bp) sequencing was performed by Macrogen, Inc. (Seoul, Korea). We generated, on average, 134,312,269 total sequence reads and approximately 71,531 Mbp of total mapped base.

### Pyrosequencing

The PyroMark PCR Kit (Qiagen, Hilden, Germany) was used for pyrosequencing with a forward primer (Pyro-FOXL2-F: 5′-AGAAGGGCTGGCAAAATAGCATC-3′) and reverse biotinylated primer (Pyro-FOXL2-R: 5′-CCGGAAGGGCCTCTTCAT-3′), or a reverse primer (Pyro-FOXL2-R2: 5′-TAGTTGCCCTTCTCGAACATGTC-3′) and a forward biotinylated primer (Pyro-FOXL2-F2: 5′-CATCGCGAAGTTCCCGTTCTA-3′). The PCR products were purified using streptavidin Sepharose HP beads (GE Healthcare, Buckinghamshire, UK), followed by hybridization with the sequencing primers (FOXL2-seqF: 5′-CGCAAGGGCAACTACT-3′ or FOXL2-seqR2: 5′-CCTTCTCGAACATGTCT-3′), as described in the PyroMark Q48 vacuum workstation guide (Qiagen). The sequencing data were analyzed using PyroMark Q48 software (Qiagen). The Pyrosequencing was performed and analyzed by Macrogen, Inc.

### Allele-specific PCR analysis

To amplify each *FOXL2* allele, allele-specific primers were designed with a 3′ mismatch at the variable nucleotide at 402. The PCR primers used for *FOXL2* amplification were the FOXL2-279F primer (5′-GAATAAGAAGGGCTGGCAAAAT-3′), the WT-specific reverse primer (5′-CCTTCTCGAACATGTCTTCG-3′), and the 402C>G-specific reverse primer (5′-CCTTCTCGAACATGTCTTCC-3′). For *GAPDH* amplification, the GAPDH-F (5′-AGGGGCCATCCACAGTCTT-3′) and GAPDH-R (5′-AGCCAAAAGGGTCATCATCTCT-3′) primers were used. PCR was performed in 20-µl reactions containing 50 ng gDNA or cDNA with DMSO (10%). PCR was performed for 34 cycles using 2 annealing temperatures: 50°C for cDNA from FFPE sections and 60°C for cDNA from cell lines. SP-Taq DNA polymerase and 2X HOT MasterMix with Dye were purchased from Cosmo Genetech and MGmed (Seoul, Korea), respectively. The PCR products were electrophoresed on a 2% agarose gel and visualized under ultraviolet (UV) light.

### Reverse transcription-quantitative real-time PCR (RT-qPCR) analysis

Following confirmation of the quality and quantity of extracted total RNA samples with a Nanodrop 2000 (Thermo Scientific, Wilmington, DE, USA), cDNA was synthesized using an iScript cDNA Synthesis Kit (Bio-Rad Laboratories, Hercules, CA, USA). Samples for RT-qPCR were prepared in a final volume of 10 µl and analyzed using Applied Biosystems’s SYBR Green PCR master mix (Bio-Rad Laboratories) during 40 cycles of amplification. RT-qPCR was performed on a CFX-96 Thermal Cycler and Detection System (Bio-Rad Laboratories). The same primers described above for the allele-specific PCR analysis were used for RT-qPCR analysis of *FOXL2* and *GAPDH* expression. Additional PCR primers used for RT-qPCR analysis were as follows: Dicer-F (5’-TCTCTTTCCCAACTGGCATC-3’), Dicer-R (5’-GGTGGTTCGTTTTGATTTGC-3’), ZEB2-F (5′-CAAGAGGCGCAAACAAGC-3′), ZEB2-R (5′-GGTTGGCAATACCGTCAT-3′), BCL-2-F (5′-GTGGAGGAGCTCTTCAGGGA-3′), BCL-2-R (5′-AGGTGCCGGTTCAGGTACTC-3′), p21-F (5′-TGTCCGTCAGAACCCATGC-3′), p21-R (5′-AAAGTCGAAGTTCCATCGCT-3′), FIS-1-F (5′-ACCTGGCCGTGGGGAACTACC-3′), FIS-1-R (5′-AGTTCCTTGGCCTGGTTGTTCTGG-3′), AFP-F (5’-CACGGATCCAACTTGAGGCTGTCATTGC-3′), AFP-R (5’-CGGAATTCGATAAGGAAATCTCACATAAAAGTC-3′), ZEB1-F (5’-ATGCAGCTGACTGTGAAGGT-3′), and ZEB1-R (5’-GAAAATGCATCTGGTGTTCC-3′). Gene-expression levels were quantified using the ΔΔCt method. All primers were purchased from Cosmo Genetech.

For miRNA analysis, ∼10 ng of total RNA was reverse transcribed using the TaqMan® microRNA Reverse Transcription Kit (Applied Biosystems, Foster City, CA, USA) and the reverse-transcription primers from the TaqMan® microRNAAssay Kit (Applied Biosystems, *hsa-miR-1236* [Assay ID: 002761]; *RNU6B* [Assay ID: 001093]). RT-qPCR was performed in a CFX-96 Thermal Cycler and Detection System, using the TaqMan® microRNA Universal PCR Master Mix (Applied Biosystems) and TaqMan probes from the TaqMan® microRNA Assay Kit, according to the manufacturer’s instructions. miRNA-expression levels were normalized to endogenous *RNU6B* expression.

### mRNA-decay rate

To assess mRNA turnover, RNA synthesis was terminated by adding 5 µg/mL ActD (Sigma-Aldrich) to the cell culture medium. At different time points (0, 0.5, 1, 2, 4, and 8 h) after ActD addition, the cells were harvested, and total RNA was isolated using the TRIzol reagent following the manufacturer’s protocol. mRNA levels were determined by real-time PCR, as described above.

### RNA interference

The target sequences of the siRNAs used in this study were as follows: siFOXL2 (5′-GGCAUCUACCAGUACAUCAdTdT-3′), siAGO1 (5′-GCACGGAAGUCCAUCUGAAUU-3′ and 5′-GAGAAGAGGUGCUCAAGAAUU-3′), siAGO2 (5′-GCACGGAAGUCCAUCUGAAUU-3′ and 5′-GCAGGACAAAGAUGUAUUAUUdTdT-3′), siAGO3 (5′-GCAUCAUUAUGCAAUAUGAUU-3′, 5′-GAAAUUAGCAGAUUGGUAAUU-3′, and 5′-CAAGAUACCUUACGCACAAUU-3′), siAGO4 (5′-GGCCAGAACUAAUAGCAAUUU3′ and 5′-GGCCAGAACUAAUAGCAAUUU-3′), siDHX9 (5′-CCAGGCAGAAATTCATGTGTG-3′ and 5′-CAAAUCAUCUGUUAAUUGUdTdT-3′), siGW182 (5′-UGAUUGUUAGGCAUCUGGCdTdT-3′ and 5′-GCCAGAUGCCUAACAAUCA-3′), siADARBP1(5’-GAUAGACACCCAAAUCGUAdTdT-3′), siDDX20 (5’-GGAAAUAAGUCAUACUUGG-3′), siFKBP5 (5’-GGAGCAACAGUAGAAAUCCdTdT-3′), siGEMIN4 (5’-GGCACUGGCAGAAUUAACA-3′), siGPKOW (5’-CUGUGUAUGUCGGACAGAUdTdT-3′), siHSP90AA1 (5’-AACAUGAAACUCAAAAAGCAUdTdT-3′), siHSP90B1 (5’-GAAGAAGCAUCUGAUUACCdTdT-3′), siLIG4 (5’-GCUAGAUGGUGAACGUAUG-3′), siMSI1 (5’-GGAGAAAGUGUGUGAAAUUdTdT-3′), siNOTCH2 (5’-GGAGGUCUCAGUGGAUAUAdTdT-3′), siPAFAH1B1 (5’-UUUAGUCUCAGAUCCUGUUGCUUCAdTdT-3′), siPPID (5’-GCAGGGAGCAAUUGACAGUdTdT-3′), and siYTHDF1 (5’-CCGCGUCUAGUUGUUCAUGAA-3′). The sequence of the control siRNA was 5′-CCUACGCCACCAAUUUCGU-3′. All siRNAs were purchased from Bioneer (Seoul, Korea). Sense and antisense oligonucleotides were annealed in the presence of annealing buffer (Bioneer).

### Immunoblot analysis

KGN cells (1 × 10^6^) and COV434 cells (0.3 × 10^6^) were transfected with the indicated plasmids or oligonucleotides using Lipofectamine 2000 (Invitrogen), according to the manufacturer’s instructions. Cell lysates were prepared and subjected to sodium dodecyl sulfate-polyacrylamide gel electrophoresis (SDS-PAGE) for subsequent immunoblotting with the respective antibodies. The protein signals on the membranes were detected using a ChemiDoc XRS + System Imager (Bio-Rad Laboratories), and the intensity of each band was quantified using Quantity One software (Bio-Rad Laboratories). For all immunoblot images presented in this manuscript, the membrane was sectioned according to the estimated molecular weights of the proteins of interest and probed with the indicated antibodies. All cropped blots were processed under the same experimental conditions. The following antibodies were used in this study: rabbit anti-FOXL2 (Park *et al*., 2010), rabbit anti-AGO1 (5053S; Cell Signaling Technology, Danvers, MA, USA), rat anti-AGO2 (11A9; Helmholtz Zentrum München, Germany), rabbit anti-AGO3 (5054S; Cell Signaling Technology), rabbit anti-AGO4 (6913S; Cell Signaling Technology), mouse anti-FLAG (F1804; Sigma-Aldrich), rabbit anti-DHX9 (ab26271; Abcam, Cambridge, MA, USA), rabbit anti-GW182 (NBP1-28751; Novus Biologicals, Littleton, CO, USA), mouse anti-Myc (2276S; Cell Signaling Technology), rabbit anti-GAPDH (sc-25778; Santa Cruz Biotechnology, Santa Cruz, CA, USA) and rabbit control-IgG (sc-2027; Santa Cruz Biotechnology).

### miRNAs and antisense miRNAs

Sense-strand RNA oligonucleotides for miR-1236 (5**′**-CCUCUUCCCCUUGUCUCUCCAG-3′), miR-1236-G (5**′**-CCUCUUCGCCUUGUCUCUCCAG-3′), miR-1236-U (5′-CCUCUUCUCCUUGUCUCUCCAG-3′), and the non-specific control RNA (5′-CCUCGUGCCGUUCCAUCAGGUAGUU-3′)(Song, Yoon *et al*., 2013) were supplied by Genolution Pharmaceuticals, Inc. (Seoul, Korea). Antisense miRNAs for mature miR-1236 (5′-CTGGAGAGACAAGGGGAAGAGG-3′), miR-145 (5′-AGGGATTCCTGGGAAAACTGGAC-3′), miR-204 (5′-AGGCATAGGATGACAAAGGGAA-3′), miR-423-3p (5′-ACTGAGGGGCCTCAGACCGAGCT-3′), and miR-484 (5′-ATCGGGAGGGGACTGAGCCTGA-3′), and a negative control miRNA (22-mer scrambled probe with a random sequence of 5′-GTGTAACACGTCTATACGCCCA-3′) (Nielsen, Jorgensen *et al*., 2011), with no known complementary sequence target among human transcripts, were purchased from Cosmo Genetech.

### Primer-extension analysis

KGN cells were transfected with the empty pCMV vector or the pCMV-FOXL2-402C>G plasmid for 48 h, and total RNA extracted using TRIzol. Total RNAs were extracted from KGN cells, COV434 cells, and AGCT tissues from three patients. Reverse transcription of total RNA was performed with poly (dT) and random hexamers (Invitrogen), according to the manufacturer’s protocol. First-strand cDNA was prepared using a forward primer (F; 5′-CGCGAGGGCGGCGGCGAGCGCAAC-3′) and a reverse primer (R; 5′-CTTCATGCGGCGGCGGCGCCGGTA-3′), and gel-purified using the QIAquick Gel Extraction Kit (Qiagen, Hilden, Germany). The reverse primer R3 (5′-GCGCCGGTAGTTGCCCTTCTC-3′) was labeled at the 5′ end using [γ-^32^P]-ATP (PerkinElmer Inc., Waltham, MA, USA) and T4 polynucleotide kinase (Takara). The extension reaction was performed with the *AccuPower*® DNA Sequencing Kit (Bioneer), and the reaction products were resolved on a 10% denaturing polyacrylamide gel and visualized by autoradiography.

### Full-length cDNA preparation using the CAGE method

Full-length cDNA was generated by the CAGE method (Takahashi, Lassmann *et al*., 2012). Briefly, 5 μg of total RNA was prepared from KGN cells. cDNA was reverse transcribed using reverse transcriptase (Takara), biotinylated with Biotin (Long Arm) Hydrazide (Vector Laboratories, Burlingame, CA, USA), and cap-trapped to capture 5′-completed cDNAs using Streptavidin C1 Dynabeads (Invitrogen).

### Genomic engineering of CRISPR/Cas9-nickase-mediated miR-1236 KO cell lines

miR-1236 KO cells were generated based on a protocol described by Ran *et al*.(Ran *et al*., 2013). Briefly, to generate vectors encoding the Cas9 (D10A) nickases, pSpCas9n(BB)-2A-GFP (Addgene, Cambridge, MA, USA) was mutagenized by PCR amplification. The primers used for site-specific mutagenesis were D10A-F (5′-TAGAGGTACCCGTTACATAAC-3′), D10A-R (5′-CTGAAGATCTCTTGCAGATAG-3′), Mut-D10A-F (5′-TACAGCATCGGCCTGGCCATCGGC-3′), and Mut-D10A-R (5′-GGTGCCGATGGCCAGGCCGATGCT-3′). Four guide RNAs were designed targeting the protospacer adjacent motif (PAM) sequence at the miR-1236* region (sgRNA-miR-1236-1 and - 2) or miR-1236 region (sgRNA-miR-1236-3 and −4) (Appendix Fig S4A and S4E), using CRISPR DESIGN (http://crispr.mit.edu/). To generate plasmids targeting miR-1236* or miR-1236, the pX458(D10A)-miR-1236-5p or pX458(D10A)-miR-1236-3p dual-guide oligonucleotide primers, respectively, were cloned into the pX458(D10A) vector. KGN or COV434 cells (1 × 10^6^) were cultured in 100-mm dishes and transfected with 5 µg of each CRISPR plasmid. The transfected cells were cultured for 24 h, harvested, and resuspended in phosphate buffered saline (PBS; Ca^2+^/Mg^2+^-free, 1 mM EDTA, 25 mM HEPES pH 7.0), containing 1% heat-inactivated FBS. They were then sorted by flow cytometry (based on green fluorescent protein signaling), using a BD FACSAria II cell sorter (BD Bioscience, San Jose, CA, USA). Cells were sorted into individual wells of 96-well plates and then further expanded.

### T7 endonuclease I (T7E1) assays

A gDNA fragment (762-bp) harboring miR-1236 was amplified with the primers, 5′-GATGGATGAAGCTTCCA-3′ and 5′-ACTCAGAATGGTACAGC-3′. The purified PCR products were denatured and reannealed in NEBuffer 2 (New England BioLabs, Ipswich, MA, USA), using a thermocycler. The hybridized PCR products were digested with T7E1 (New England BioLabs) for 2 h and resolved on 2% agarose gels.

### Genotyping for miR-1236 KO

Allelic deletion was confirmed using the TOPcloner™ TA Core Kit (Enzynomics, Daejeon, Korea) and DNA sequencing (Cosmo Genetech).

### Northern blot analysis

Total RNA (including miRNAs) was extracted from cultured cells using TRIzol (Invitrogen). Then, the small RNA species were isolated and concentrated using the mirVana PARIS Kit (Ambion). Small RNA fractions from cells (∼1.5 μg) were separated on a 15 % polyacrylamide gel containing 8 M urea, visualized by ethidium bromide staining, and then transferred to a Hybond-XL membrane (Amersham Bioscience). Subsequently, the RNA was immobilized on the membrane with a ultraviolet cross-linker (UVP) and hybridized with a 5′ end-labeled-oligonucleotide. Anti-miR-1236 (5′-CTGGAGAGACAAGGGGAAGAGG-3′) and anti-miR-21 (5′-TCAACATCAGTCTGATAAGCTA-3′) were used as probes. The radioactive signals were analyzed using a Bio-Rad phosphorimager and the Quantity One image-analysis software package (Bio-Rad).

### Luciferase-reporter assays

Luciferase assays were performed as described previously (Kim *et al*., 2011).

### *In vitro* transcription

*In vitro* transcription of 230-bp CDS sequences that included nucleotide 402 of WT and 402C>G *FOXL2* mRNA was performed using the MegashortScript^TM^ T7 Kit (Thermo Scientific, Rockford, IL, USA). To generate the T7-FOXL2 DNA template, pCMV-FOXL2-WT and pCMV-FOXL2-C134W (Kim *et al*., 2011) were used as templates and amplified by PCR, with a primer containing the T7 promoter sequence, T7-FOXL2-F (5′-TAATACGACTCACTATAGGGAGGGCTGGCAAAATAGCA-3′) and T7-FOXL2-R (5′-GGCGCCTCCGGCCCCGA-3′). RNA was transcribed *in vitro* using T7 RNA polymerase according to the manufacturer’s instructions.

### *In vitro*-annealing kinetics between miR-1236 and *FOXL2* transcripts

We incubated 5′-labeled ^32^P-miR-1236 (0.5 nM; PerkinElmer, Boston, MA, USA) with increasing concentrations of *in vitro-*transcribed WT or 402C>G *FOXL2* mRNAs (0, 2.5, 12.5, 25, and 50 nM) in hybridization buffer (100 mM NaCl, 20 mM Tris-HCl at pH 7.4, and 10 mM MgCl_2_) at 65°C for 5 min. Then, the hybridization reactions were cooled slowly to room temperature. Aliquots were transferred into 1 volume (v) of stop buffer (20 mM Tris-HCl at pH 7.4, 10 mM EDTA, 2% [v/v] SDS, 8 M urea, 0.025% [v/v] bromophenol blue) and analyzed by 6% native PAGE. Band intensities reflecting *FOXL2*–miRNA complex formation were detected and quantified using a phosphorimager and the OptiQuant software package (Packard, Meriden, CT, USA).

### *In vivo* association of *FOXL2* mRNAs or miR-1236 with RISC

293T cells were cotransfected with miR-1236-G mimic, siDHX9, or the indicated plasmids for 48 h. The cells were washed with cold 1× PBS, and RNA–protein complexes were cross-linked by UV-A irradiation (0.15 J/cm^2^). In each case, 10% of the cells was removed and kept as the input samples, and remaining cells were lysed in buffer containing 25 mM Tris-HCl at pH 7.4, 150 mM KCl, 0.5% NP-40, 2 mM EDTA, 1 mM NaF, 0.5 mM DTT, RNase inhibitor (Promega), 0.05% Tween 20, and protease inhibitors (GeneDEPOT, Barker, TX, USA), and centrifuged at 7,000 × *g* for 15 min at 4°C. Each lysate was pre-incubated with 2 μg IgG and 20 μl Dynabeads (Invitrogen) for 4 h at 4°C. Three milligrams of each lysate was precleared using a control rabbit IgG-Dynabeads complex for 30 min at 4°C. Then, the cell lysates were re-incubated with 5 μg FLAG antibody-coupled Dynabeads (50 μl) overnight on a rotator at 4°C. After incubation, the beads were washed five times with a buffer (300 mM KCl, 50 mM Tris-HCl pH 7.4, 1 mM MgCl_2_, 0.1% NP-40, 0.05% Tween 20) and incubated with DNase I for 10 min at 37°C. The sample proteins were digested with proteinase K, and RNA was extracted using an acidic phenol: chloroform mixture (5: 1, pH 4.3) and precipitated with isopropanol in the presence of 10% of 3 M sodium acetate (NaOAc; pH 5.2). The purified RNAs were reverse transcribed to cDNA with the FOXL2-430-R primer (5′-GGTAGTTGCCCTTCTCGAAC-3′), which binds to a region downstream of the 402C>G site. Then, the cDNA products were used for *FOXL2* allele-specific RT-PCR or real-time RT-PCR analysis, with the FOXL2-279F primer. The relative abundance of miR-1236 was validated using TaqMan® microRNA assays. To detect endogenous association of AGO3-mediated RISCs and *FOXL2* mRNA, RNA–protein complexes formed in KGN cells (3 × 10^7^) transfected with a control miRNA or miR-1236-G (10 nM), or a control siRNA or siDHX9 (200 nM), were cross-linked via UV irradiation (0.15 J/cm^2^). The lysates were incubated with 10 μl of AGO3 antibody-coupled Dynabeads (50 μl) overnight on a rotator at 4°C. The RNAs were purified as described above.

### Cell-viability assay

Cell-viability assays were performed as previously described (Kim *et al*., 2014).

### 5-bromo-2′-deoxy-uridine cell-proliferation assay

Cell (1 × 10^4^) proliferation was measured using the 5-Bromo-2′-deoxy-uridine Labeling and Detection Kit III (Roche, Mannheim, Germany) (Jin, Lee *et al*., 2016).

### Cell-migration assay

A chemotaxis migration assay was performed using Transwell Permeable Supports (8-μm pore size, 6.5-mm insert; Corning-Costar, Lowell, MA, USA). KGN cells (1 × 10^6^) and COV434 cells (0.3 × 10^6^) were transfected with either pcDNA3 or pcDNA3-Flag-FOXL2)(Park *et al*., 2010), or the indicated oligonucleotides. Cells were plated at a density of 2 × 10^5^ cells/well in the top chamber containing DMEM/F12 and 1% FBS, whereas the bottom chamber contained DMEM/F12 and 10% FBS as a chemoattractant. After 2 to 6 h of incubation, the cells and medium were removed from the top chamber, and the migrated cells were fixed with 5% glutaraldehyde for 20 min and stained with 0.5% crystal violet for 20 min. Images of the migrated cells were taken at ×100 magnification under a bright-field microscope. Quantification was performed under a light microscope with ×200 magnification by counting the number of migrated cells in five random fields per chamber. The data are presented as the mean ± SEM of three independent experiments and are shown as the fold-change in the average number of cells.

### Apoptosis and cell-cycle analyses

Detection of Annexin V-positive apoptotic cells and cell-cycle analysis were performed as previously reported (Jin *et al*., 2016, Kim *et al*., 2011).

### Tumor xenograft establishment and immunohistochemistry

Control or miR-1236-KO (miR-1236^−/−^) KGN cells (5 × 10^6^) were suspended in 0.1 mL Matrigel (1: 1, v/v, Corning, Tewksbury, MA) and injected subcutaneously into 7-week-old BALB/c nu/nu immunodeficient mice (weights ranged between 18 and 20 g) (ORIENT BIO, Seongnam, Korea). Mice were euthanized 2 months later, and the numbers of intraperitoneal nodules were counted. Half of the nodules were excised, fixed in 4% neutral buffered formalin, and embedded in paraffin before preparing 5-μm sections that were stained with hematoxylin. The other half of the nodules were excised, and total RNA was isolated using the TRIzol reagent (Invitrogen), according to the manufacturer’s instructions. The animal guidelines were approved by the Chung-Ang University Institutional Animal Care and Use Committee (IRB# CAU2012-0044), and the animals were treated as described in the protocol.

### Statistical analysis

All details of the statistical analysis used in the experiments are included in the figure legends. Multiple-comparison analyses of values were performed using the Student–Newman–Keuls test, and Student’s *t*-test was used for comparisons with control samples, using SAS version 9.2 (SAS Institute, Cary, NC, USA) and SigmaPlot (Systat Software, San Jose, CA, USA). The data are presented as mean ± SEM, and *P* < 0.05 was considered to reflect as statistically significant difference.

### Data availability

Ultra-deep RNA Sequencing data have been deposited in the NCBI Sequence Read Archive (SRA) with the BioProject code “PRJNA555182”.

## ACKNOWLEDGEMENTS

We thank Drs. Wooseok Song, Hong-Man Kim, and Seunghwa Lee for helping with the primer-extension and cell-migration experiments. We also thank Dr. Jae-Sung Woo for critical comments. This research was supported by the National Research Foundation of Korea (NRF), which is funded by the Ministry of Science, ICT, and Future Planning (2017R1A2B2011248; 2015R1A5A1008958; 2018R1A5A1025077; 2019R1H1A2080015; 2019R1A2C2086590).

## AUTHOR CONTRIBUTIONS

K.L. and J.B. conceived and designed the study; K.L. and J.B. developed the methods; E.S., H.J., D.S.S., Y.L., and T.H.K. performed the experiments; Y.H., S.H., K.L., and J.B. analyzed and interpreted the data; E.S., H.J., K.L., and J.B. wrote and reviewed the manuscript; K.L. and J.B. supervised the study.

