## Supplemental Figures for "An alternative miRISC targeting a coding mutation site in *FOXL2* links to granulosa cell tumor"

### Appendix Figure S1

**A**

Sequence to analyze:
GGACGCTGGACCCGGCCTGC/GGAAGACATGTTCGAGAAGGGCAACTACCGGCGCCG

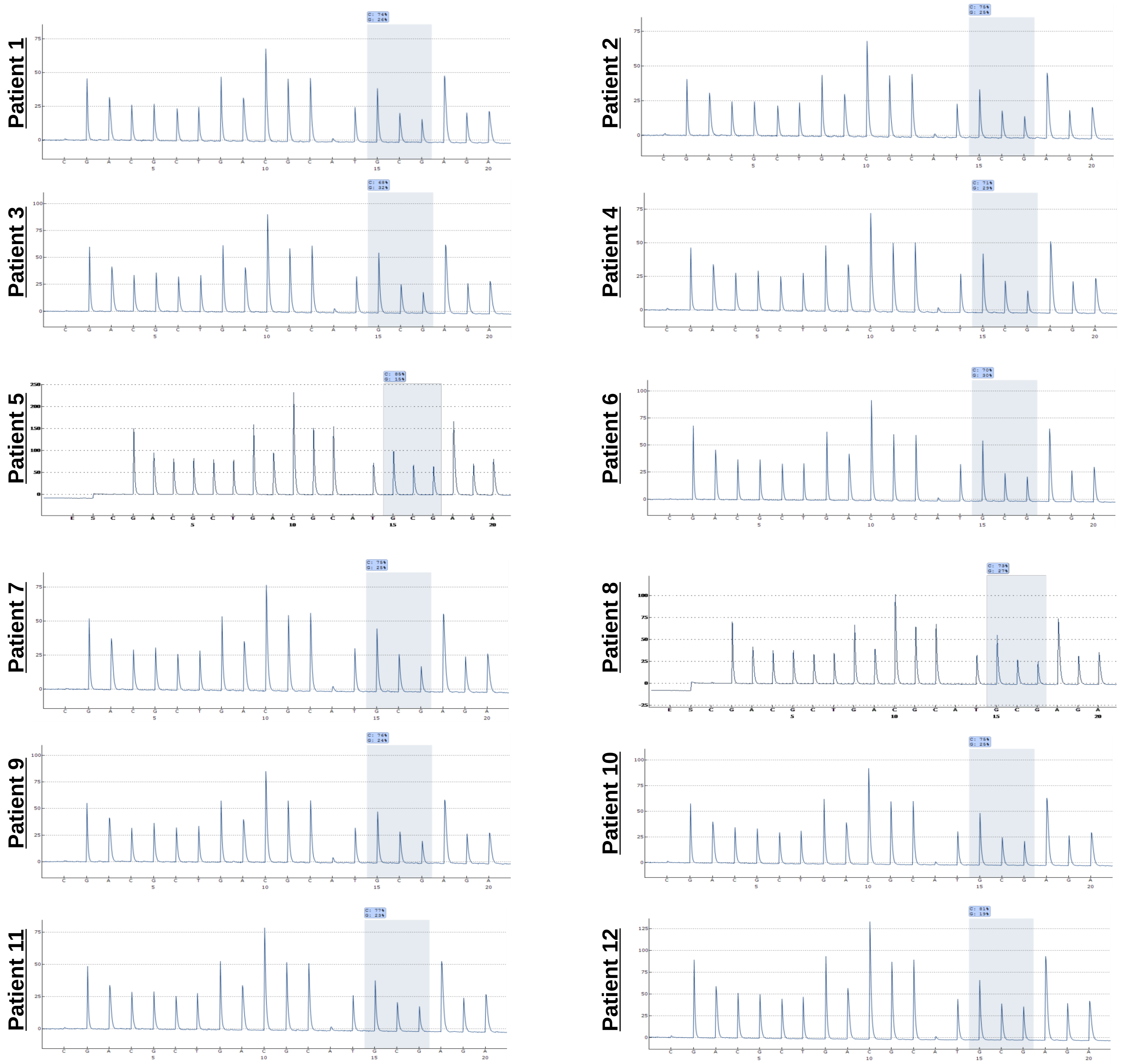


**B**

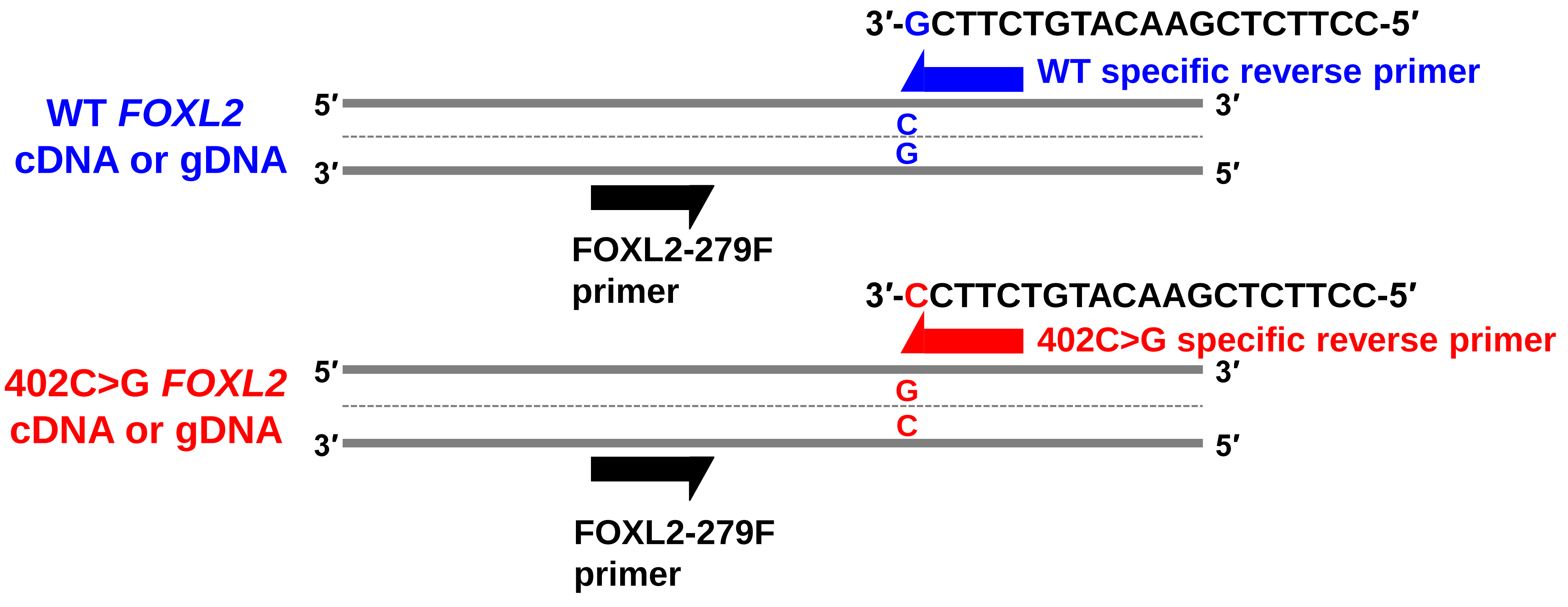

### Appendix Figure S1

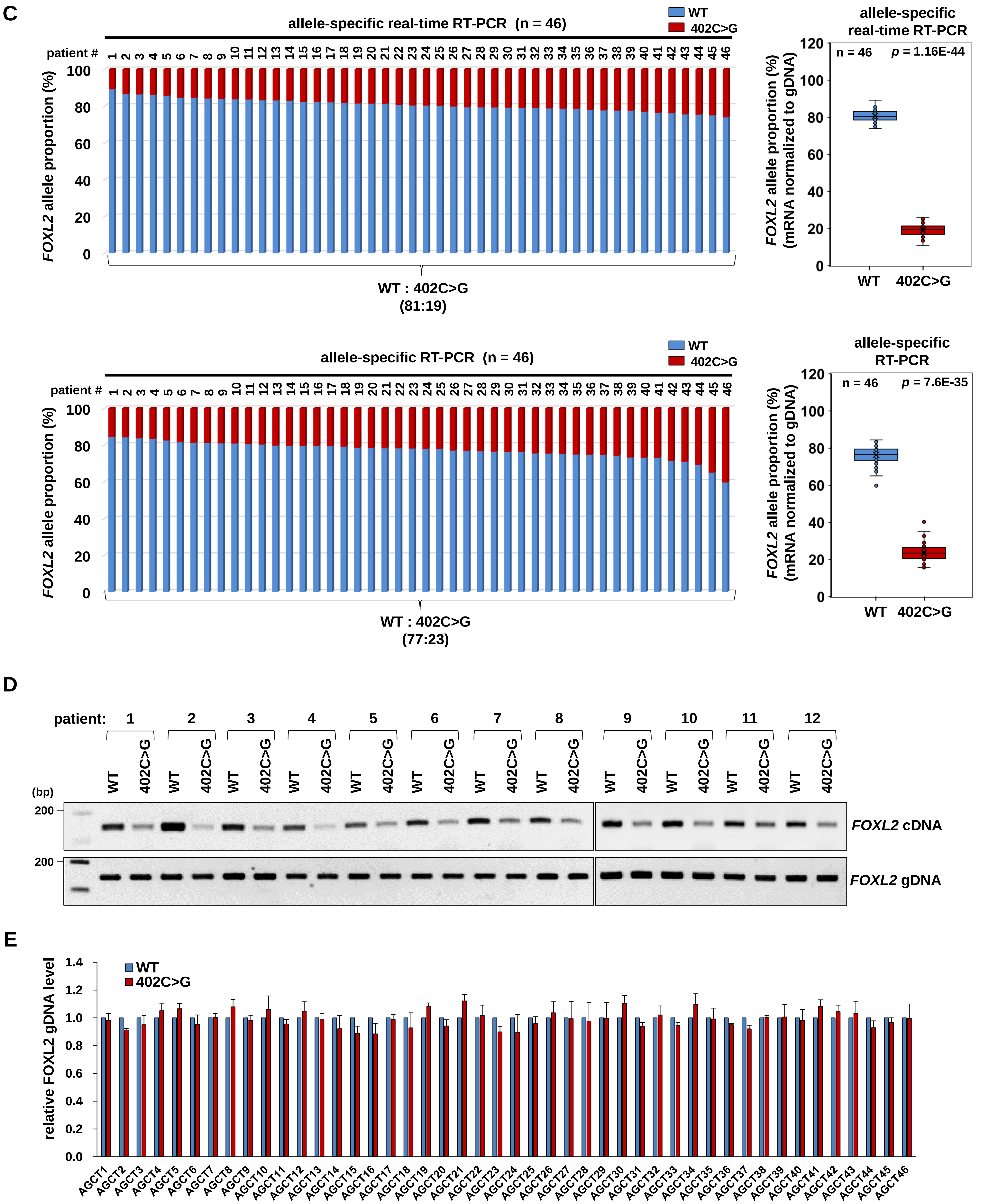

F

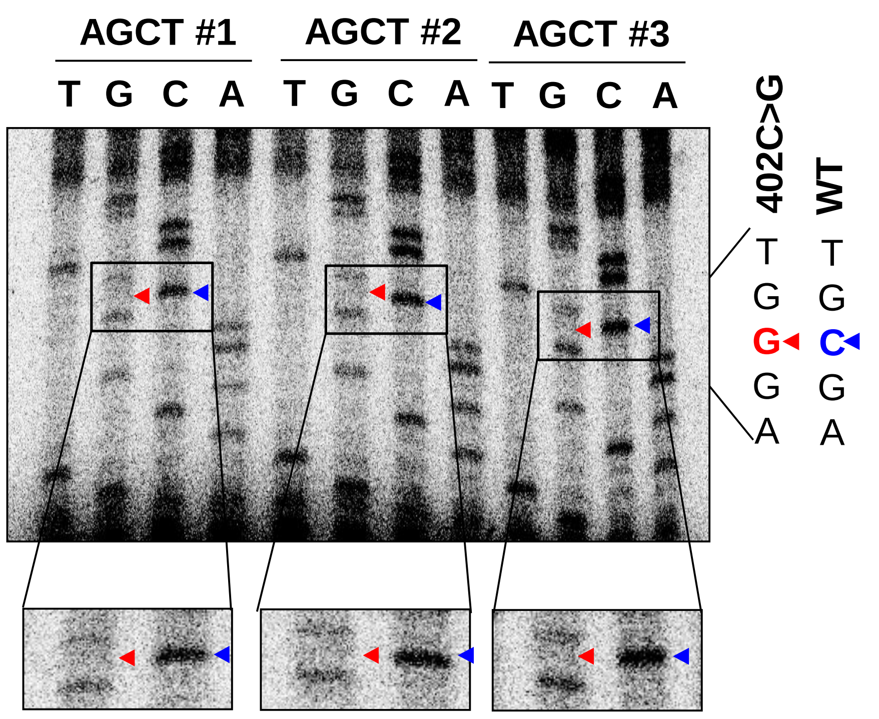
AGCT #1
AGCT #2
AGCT #3
T G C A T G C A T G C A
402C>G
WT
T G G G A
T G C G A

G

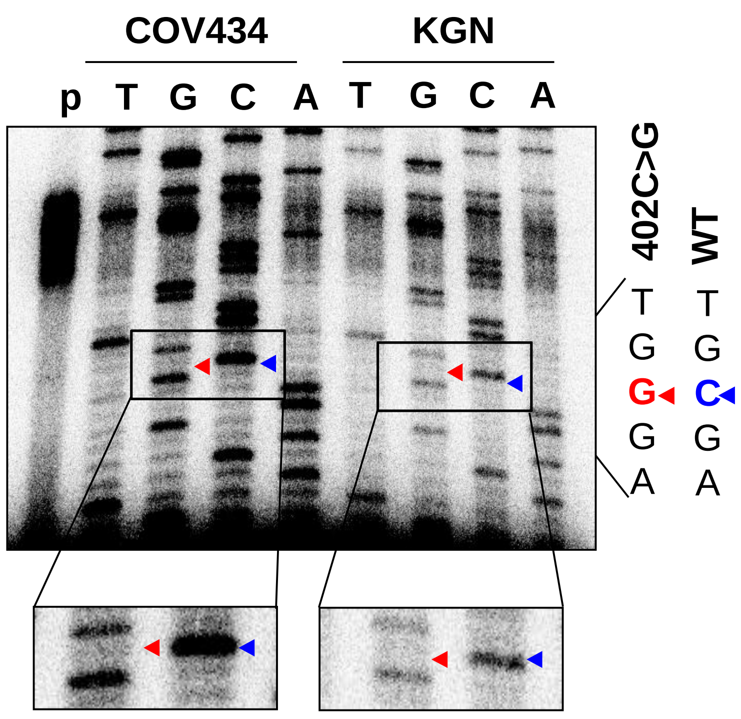
COV434
KGN
p T G C A T G C A
402C>G
WT
T G G G A
T G C G A

H

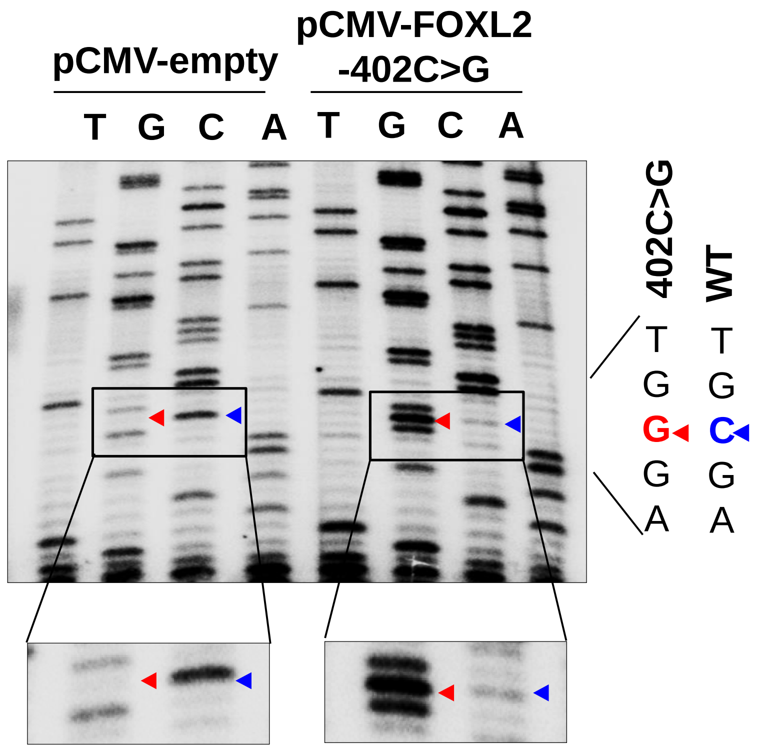
pCMV-empty
pCMV-FOXL2
-402C>G
T G C A T G C A
402C>G
WT
T G G G A
T G C G A

I

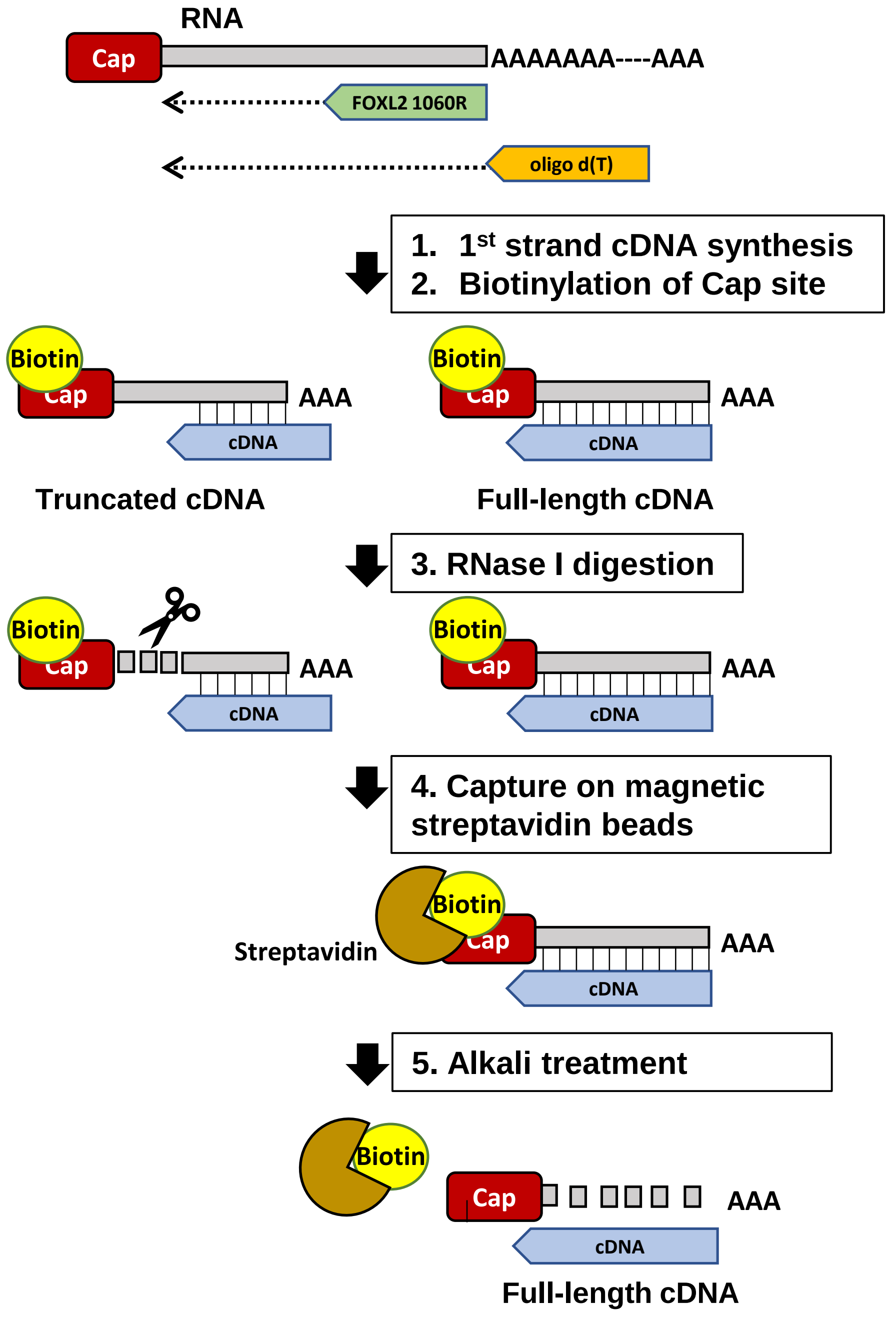
RNA
Cap
AAAAAAA----AAA
FOXL2 1060R
oligo d(T)
1. 1st strand cDNA synthesis
2. Biotinylation of Cap site
Biotin
Cap
AAA
cDNA
Truncated cDNA
Full-length cDNA
3. RNase I digestion
4. Capture on magnetic streptavidin beads
Streptavidin
5. Alkali treatment
Full-length cDNA

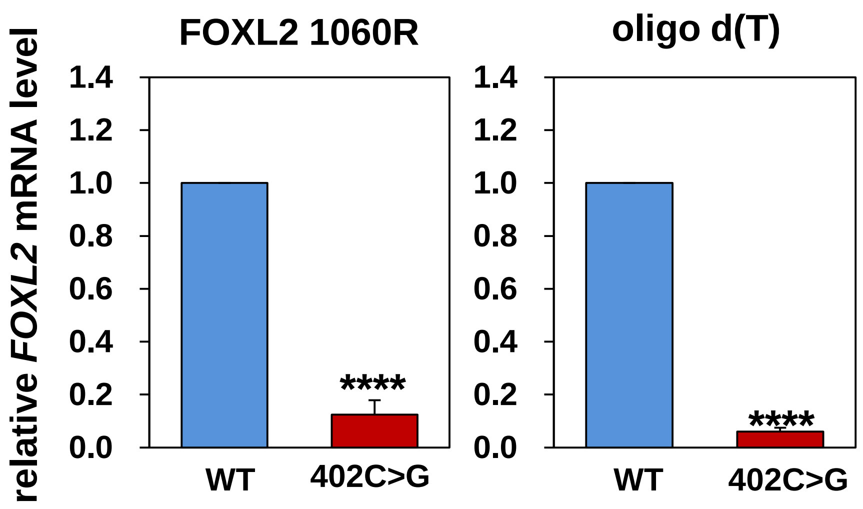
FOXL2 1060R
oligo d(T)
relative FOXL2 mRNA level
1.4
1.2
1.0
0.8
0.6
0.4
0.2
0.0
****
WT
402C>G

**Appendix Figure S1. Allelic imbalance of heterozygous *FOXL2* transcripts.**

**A** Pyrograms of WT and variant *FOXL2* mRNAs in 12 representative AGCTs analysed by pyrosequencing analysis.

**B** Schematic representation of allele-specific PCR analysis, performed with the allele-specific primers for WT and 402C>G mutant mRNAs, used in the experiments represented in Figure 1. The allele-specific PCR was based on the inability of Taq DNA polymerase to extend primers when their 3′-end nucleotides do not base-pair with template DNA. The amounts of WT and 402C>G variant *FOXL2* amplicons generated with cDNA or gDNA from KGN and COV434 cells were determined by PCR, using a common forward primer (FOXL2-279F) and allele-specific reverse primers.

**C** Bar graph and box-and-whisker plots showing the allelic proportions of WT *FOXL2* mRNA and 402C>G *FOXL2* mRNA in AGCT tissues from independent patients (allele-specific real-time RT-PCR and allele-specific RT-PCR, n = 46). Allele-specific semi-quantitative and real-time RT-PCR data were normalized to the paired-gDNA levels in each sample. The box plot represents the lower, median, and upper quartiles, and the whiskers represent the 95% confidence interval of the mean. The whiskers extend to the most extreme data points not considered outliers, and the outliers are represented as dots. The allele-specific semi-quantitative and real-time RT-PCR data are presented as the mean ± SEM from three independent experiments.

**D** The electrophoretic results of WT and variant *FOXL2* mRNAs in specimens from patients with AGCT. The relative abundances of WT and variant *FOXL2* mRNAs were analyzed in AGCT tissues from 12 patients by allele-specific RT-PCR analysis. Representative gel images are shown.

**E** The relative abundances of WT and variant *FOXL2* gDNAs isolated and analysed in AGCT tissues from 46 patients by allele-specific real-time RT-PCR. The relative abundances of the variant FOXL2 gDNA were normalized to that of WT allele (set to 1). The data (mean ± SEM) are from three independent experiments.

**F, G, H**, Primer extension results for the 402C>G locus of FOXL2 mRNAs. The relative abundance of WT and mutated transcripts of FOXL2 was analysed in COV434 or KGN cells (**F**) and AGCT tissues from three patients (**G**) by primer extension analysis as described below. (**H**) Primer extension analysis performed on total RNA extracted from KGN cells transfected with pCMV-FOXL2-402C>G or pCMV empty vector. The expected position of the 402C>G variant nucleotide is indicated by the arrowhead. In lane P, primers were loaded. The WT (C) or mutated (G) nucleotide at position 402 is indicated with bold blue or red text, respectively, throughout this paper. Reverse transcription of total RNA was performed with poly(dT) and random hexamers according to the manufacturer's protocol (Invitrogen, Carlsbad, CA, USA). First-strand cDNA was amplified with the forward primer F (5′-CGCGAGGGCGGCGGCGAGCGCAAC-3′) and the reverse primer R (5′-CTTCATGCGGCGGCGGCGCCGGTA-3′) and purified using the QIAquick gel extraction kit (Qiagen, Hilden, Germany). The reverse primer R3 (5′- GCGCCGGTAGTTGCCCTTCTC-3′) was labelled at the 5′ end using [γ-32P] ATP (PerkinElmer Inc., Waltham, MA, USA) and T4 polynucleotide kinase (TaKaRa Bio, Kyoto, Japan). The extension reaction was performed with the AccuPower® DNA Sequencing kit (Bioneer, Daejeon, Korea), and reaction products were resolved on a 10% denaturing polyacrylamide gel and visualized by autoradiography. Signals of the 402 regions of FOXL2 transcripts are enlarged under the main image.

**I** Strategies for amplifying capped full-length WT and 402C>G variant FOXL2 using the cap analysis of gene expression (CAGE) method with two different reverse primers (left). Relative abundances of WT and variant FOXL2 mRNAs were analysed in KGN cells via allele-specific real-time RT-PCR analysis with two different reverse primers (right). The data (mean ± SEM) are from three independent experiments. ****$p < 0.0001$

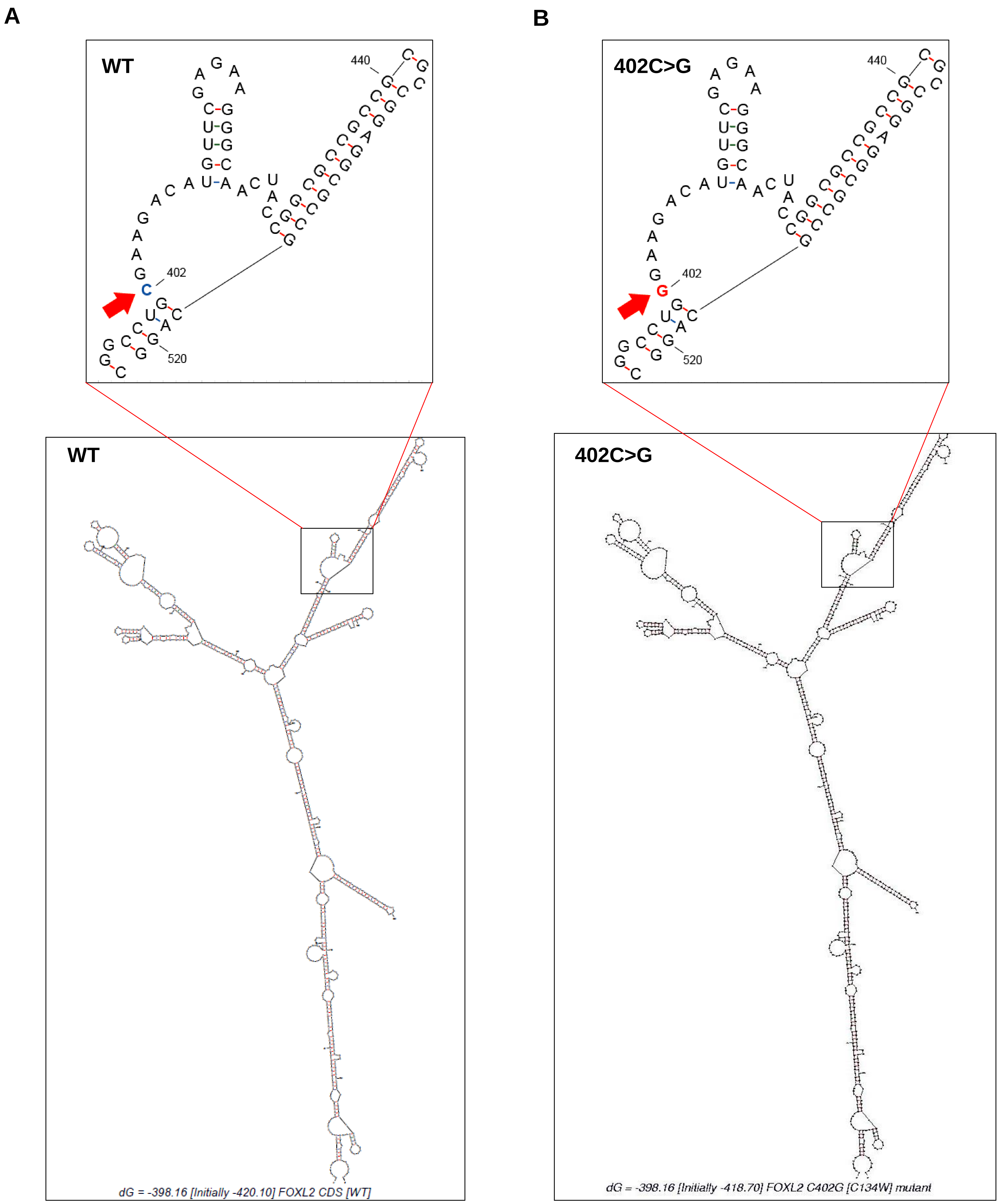


**Appendix Figure S2. Predicted secondary structures of WT and 402C>G *FOXL2* mRNAs.**

**A, B** The M-fold program (http://mfold.rna.albany.edu/?q=mfold/RNA-Folding-Form) was used to compare the secondary structures of WT (**A**) and variant (**B**) *FOXL2* mRNA. The structure in the boxed region, containing the 402C>G nucleotide change, is enlarged in the bottom panel, where 402G is indicated in red color with a red arrow.

### Appendix Figure S3

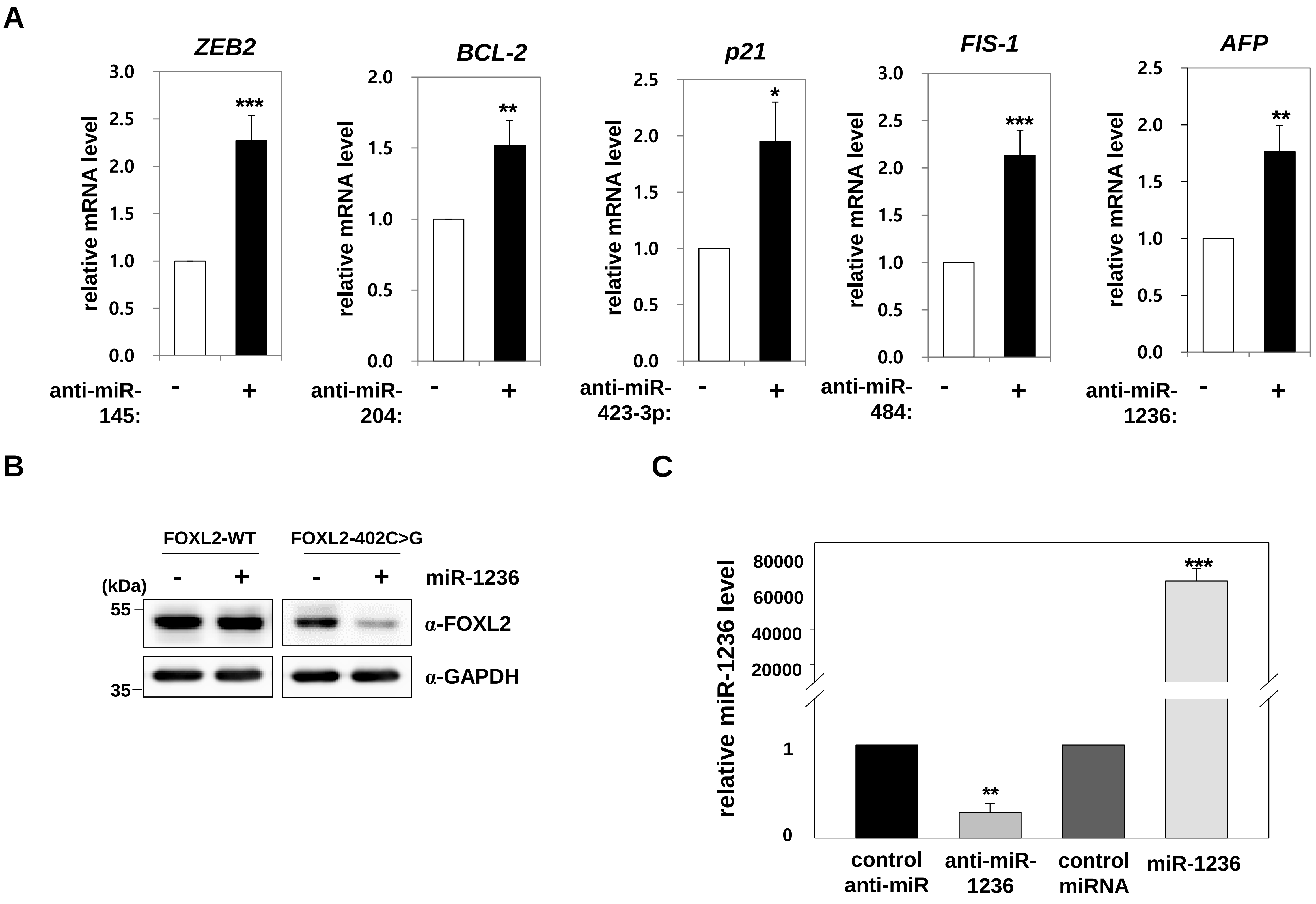


**Appendix Figure S3. Effectiveness of miRNAs and anti-miRNAs.**

**A** To validate the effectiveness of the anti-miRNAs used in Fig. 2A and B, KGN cells were transfected with anti-miR-145, anti-miR-204, anti-miR-423-3p, anti-miR-484, or anti-miR-1236 for 48 h, and the corresponding target-mRNA levels were determined by real-time RT-PCR. The data (mean ± SEM) are from three independent experiments, performed in triplicate.

**B** 293T cells were cotransfected with a control miRNA or miR-1236 mimic, and with an expression vector encoding WT or variant FOXL2. FOXL2 protein expression was assessed by western blot analysis.

**C** miR-1236 levels in KGN cells were quantitated at 48 h post-transfection with a control anti-miR, anti-miR-1236, a control miRNA, or miR-1236, using the TaqMan miRNAs Real-Time PCR Quantitation Assay Kit. The data (mean ± SEM) are from three independent experiments performed. *$p < 0.05$, **$p < 0.01$, ***$p < 0.001$

### Appendix Figure S4

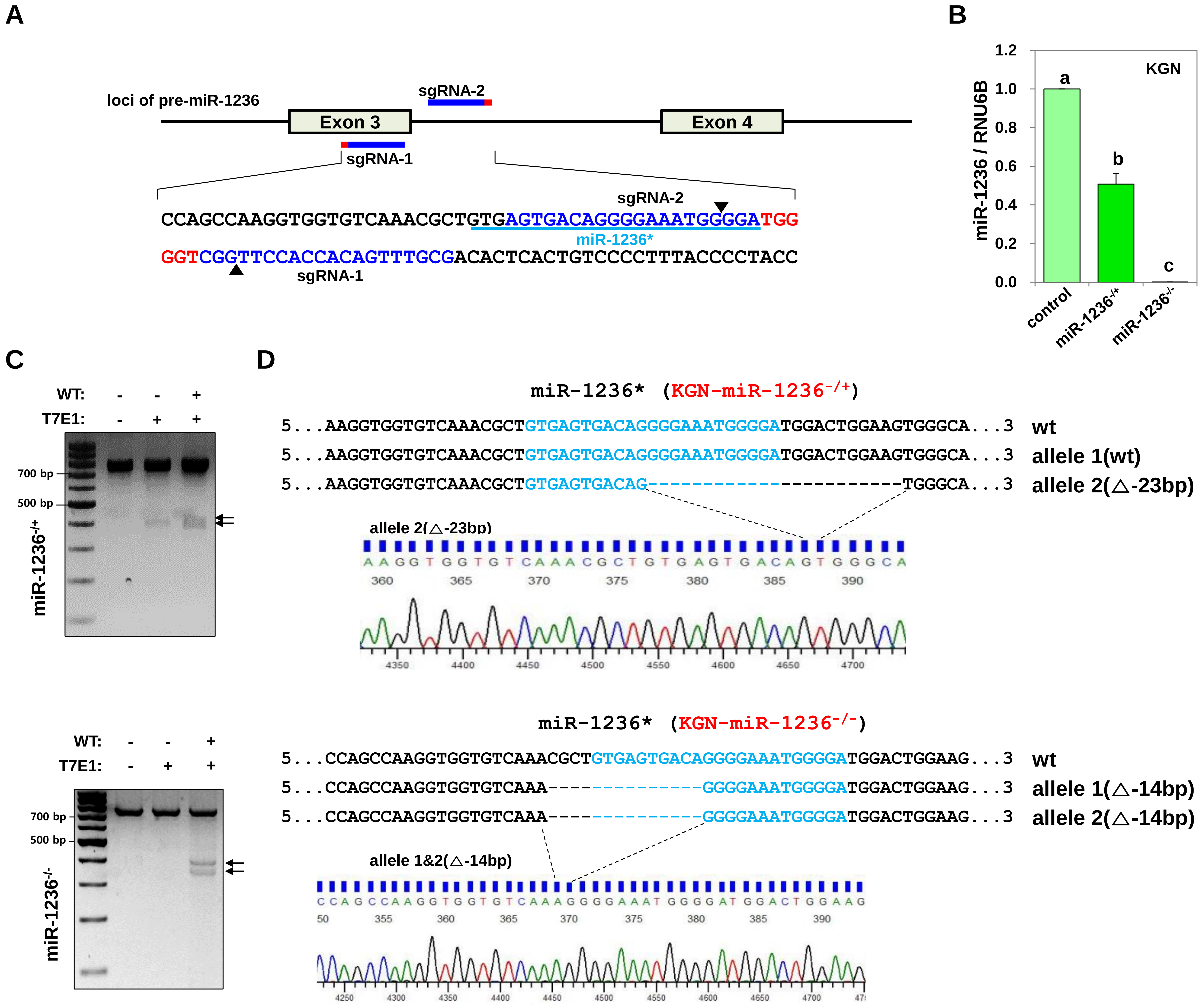

### Appendix Figure S4

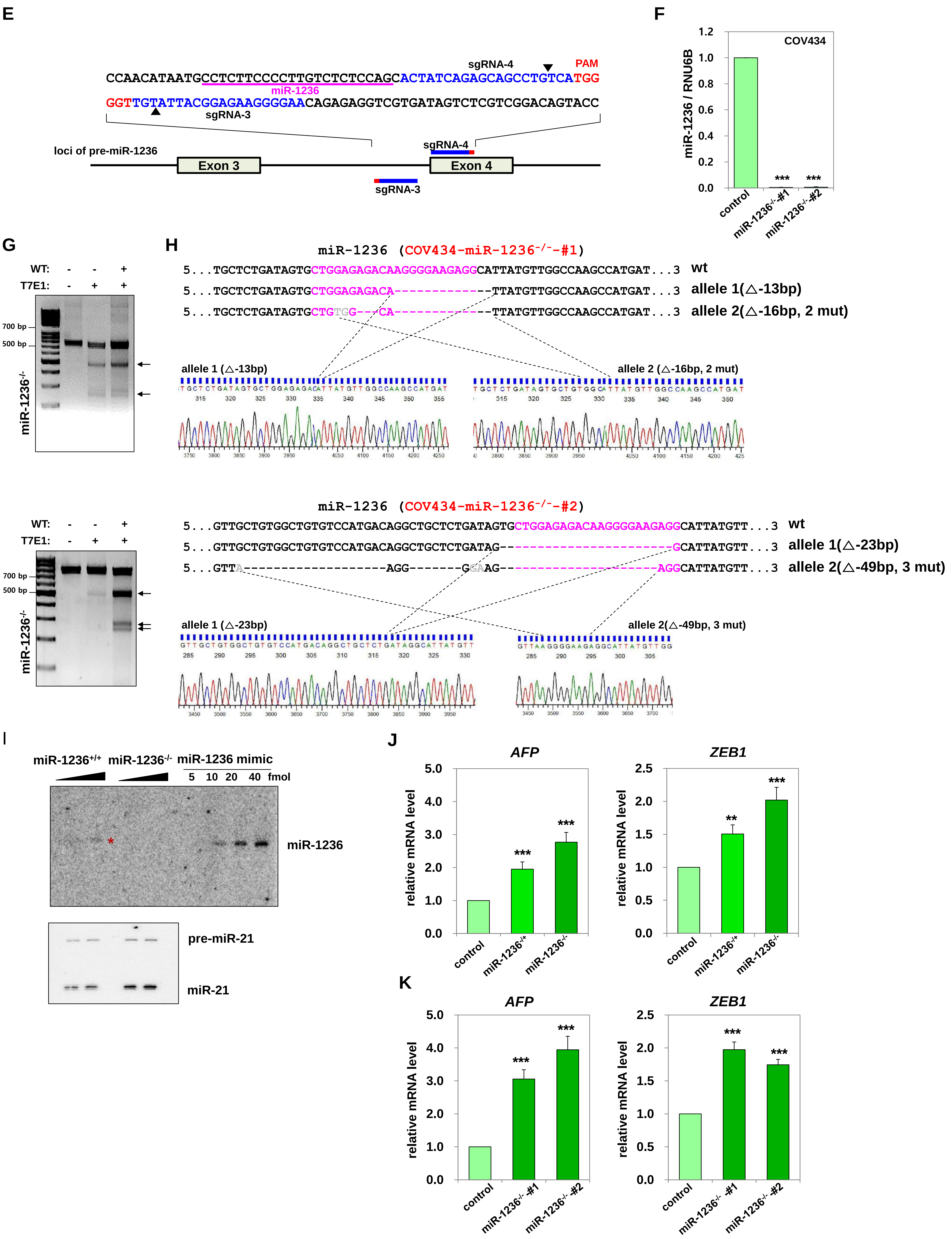

**Appendix Figure S4. *miR-1236* KO in KGN cells and COV434 cells using the CRISPR/Cas9 system.**

**A** Schematic representation of the CRISPR/Cas9-nickase strategy for *miR-1236* KO in KGN cells. A pair of single guide RNAs (sgRNA-1 and -2) was designed targeting the PAM sequence at the *miR-1236** region of pre-*miR-1236*. The mature sequence of *miR-1236** is underlined. The sgRNA and PAM sequences are shown in blue and red, respectively. The triangle indicates the possible cleavage site.

**B** KO of *miR-1236* was validated using TaqMan® microRNA assays. The relative expression level of *miR-1236* was normalized to *RNU6B* small nuclear RNA and compared with the corresponding levels in the WT cell line. The data (mean ± SEM) are from three independent experiments. Different letters denote statistically significant differences ($p < 0.001$).

**C** Results from the T7E1 assay, performed to detect CRISPR/Cas9-induced modification in *miR-1236*$^{-/+}$ KO KGN (top) and *miR-1236*$^{-/-}$ KO KGN (bottom) cells. The arrows indicate the position of the expected DNA bands, following cleavage by T7E1.

**D** DNA sequences of the target sites in the *miR-1236*$^{-/+}$ or *miR-1236*$^{-/-}$ KGN cells are presented. The *miR-1236** sequences are shown in light blue. The numbers of mutated nucleotides are indicated at the right. Sequencing chromatograms are shown.

**E** A schematic representation of the CRISPR/Cas9-nickase strategy for *miR-1236* KO in COV434 cells. A pair of sgRNAs (sgRNA-3 and -4) was designed to target PAM sequences in the *miR-1236* region of pre-*miR-1236*. The mature *miR-1236* sequence is underlined. The sgRNA and PAM sequences are shown in blue and red, respectively. The triangles indicate the possible cleavage sites.

**F** KO of *miR-1236* was validated using TaqMan® microRNA assays. The relative expression levels of *miR-1236* were normalized to that of *RNU6B* small nuclear RNA and compared with the control (WT) cell line. The data (mean ± SEM) are from three independent experiments (***$p < 0.001$).

**G** Results from T7E1 assays, performed to detect CRISPR/Cas9-induced modification in COV434 cells. The arrows indicate the position of the expected DNA bands, following cleavage by T7E1.

**H** DNA sequences of the target sites in *miR-1236*$^{-/-}$ COV434 cells are presented. The *miR-1236* sequences are shown in pink. The numbers of mutated nucleotides are indicated at the right. Sequencing chromatograms are shown.

**I** Northern blot analysis of miR-1236 in *miR-1236*$^{+/+}$ or *miR-1236*$^{-/-}$ KGN cells. Identical blots were stripped and re-probed for miR-21 analysis. The red asterisk denotes the miR-1236 band. Lanes 1–2 and 3–4 were loaded with small RNAs isolated from *miR-1236*$^{+/+}$ and *miR-1236*$^{-/-}$ cells, respectively. Lanes 1 and 3, and 2 and 4 were loaded with small RNAs isolated from $10^7$ and $2 \times 10^7$ cells, respectively.

**J, K** Increased expression of known target genes of miR-1236 in miR-1236 KO cells is shown. Changes in the levels of *AFP* and *ZEB1* mRNAs between WT and miR-1236 KO cells were determined by real-time RT-PCR in KGN (J) and COV434 (K) cells. The data (mean ± SEM) are from three independent experiments. **$p < 0.01$, ***$p < 0.001$

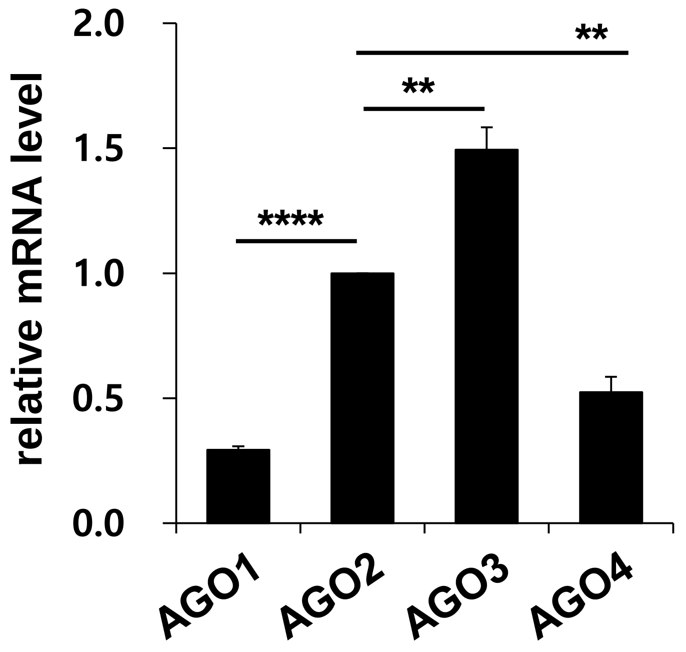


**Appendix Figure S5. Predominant expression of *AGO3* mRNA in KGN cells.**

The mRNA levels of *AGO1*, *AGO2*, *AGO3*, and *AGO4* in KGN cells were determined by real-time RT-PCR. The relative abundances of *AGO1–4* mRNA were quantified by setting the amount of *AGO2* mRNA to 1. Expression levels of *AGO1–4* mRNAs observed by real-time RT-PCR analyses were normalized to *GAPDH* mRNA levels. The data (mean ± SEM) are from three independent experiments. (**$p < 0.01$, ****$p < 0.0001$).

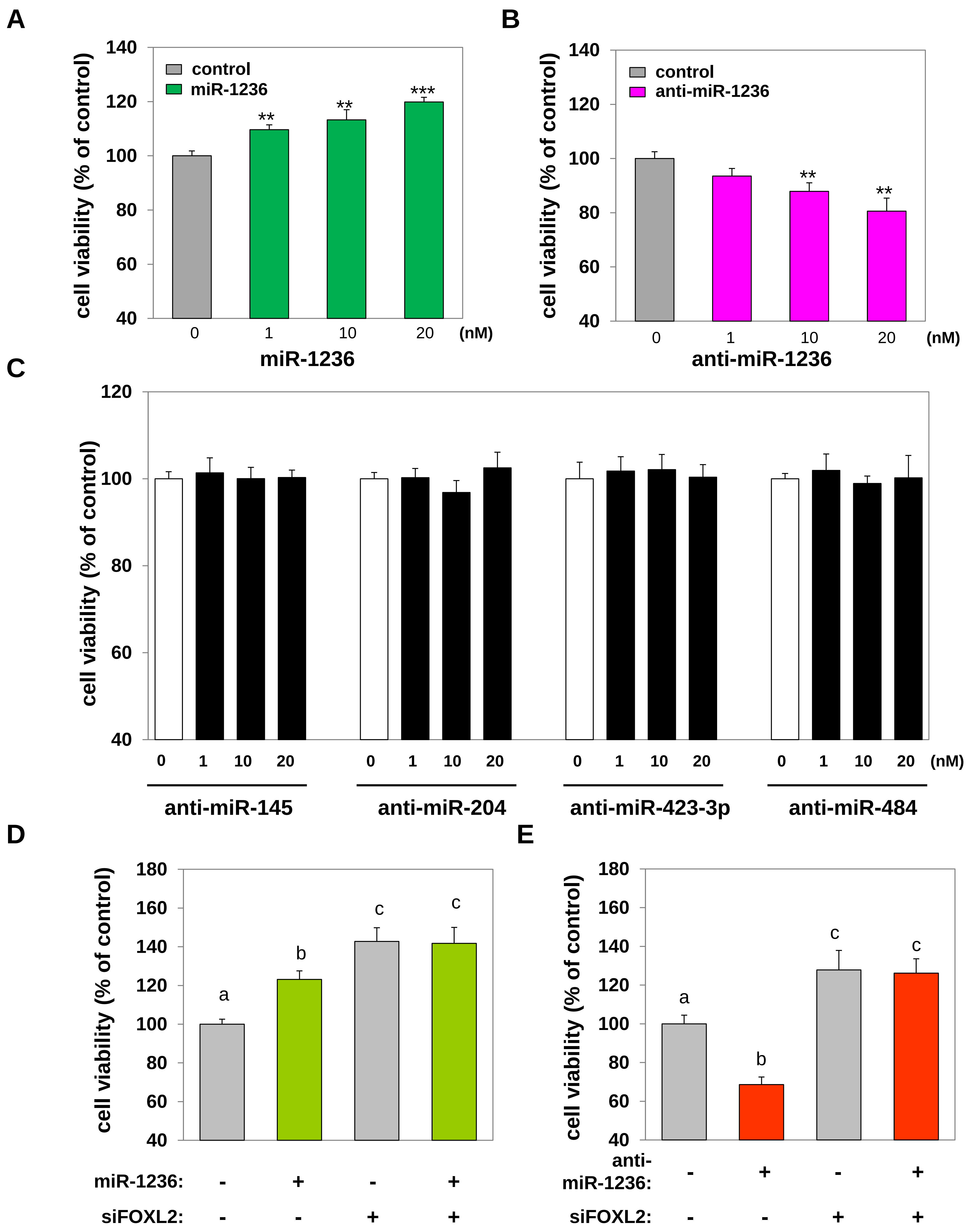


**Appendix Figure S6. Effects of miR-1236 on cell viability.**

**A, B** The effects of miR-1236 on the viability of KGN cells were assessed at 48 h post-transfection with the indicated concentrations of miR-1236 (**A**) or anti-miR-1236 (**B**). $^{**}p < 0.01$, $^{***}p < 0.001$.

**C** The activities of other candidate miRNAs on the viability of KGN cells were assessed. KGN cells were transfected with 0, 1, 10, or 20 nM of anti-miR-145, anti-miR-204, anti-miR-423-3p, or anti-miR-484 for 48 h. No statistically significant effect was observed. The data are presented as the mean ± SEM from three independent experiments, performed in triplicate.

**D, E** KGN cells were transfected with scrambled control or FOXL2-specific siRNAs (200 nM) for 24 h. These cells were treated with miR-1236 (**D**) or anti-miR-1236 (**E**) for 48 h, and cell viabilities were measured. Different letters denote statistically significant differences ($p < 0.0001$). The data presented are the mean ± SEM of three independent experiments, performed in triplicate.

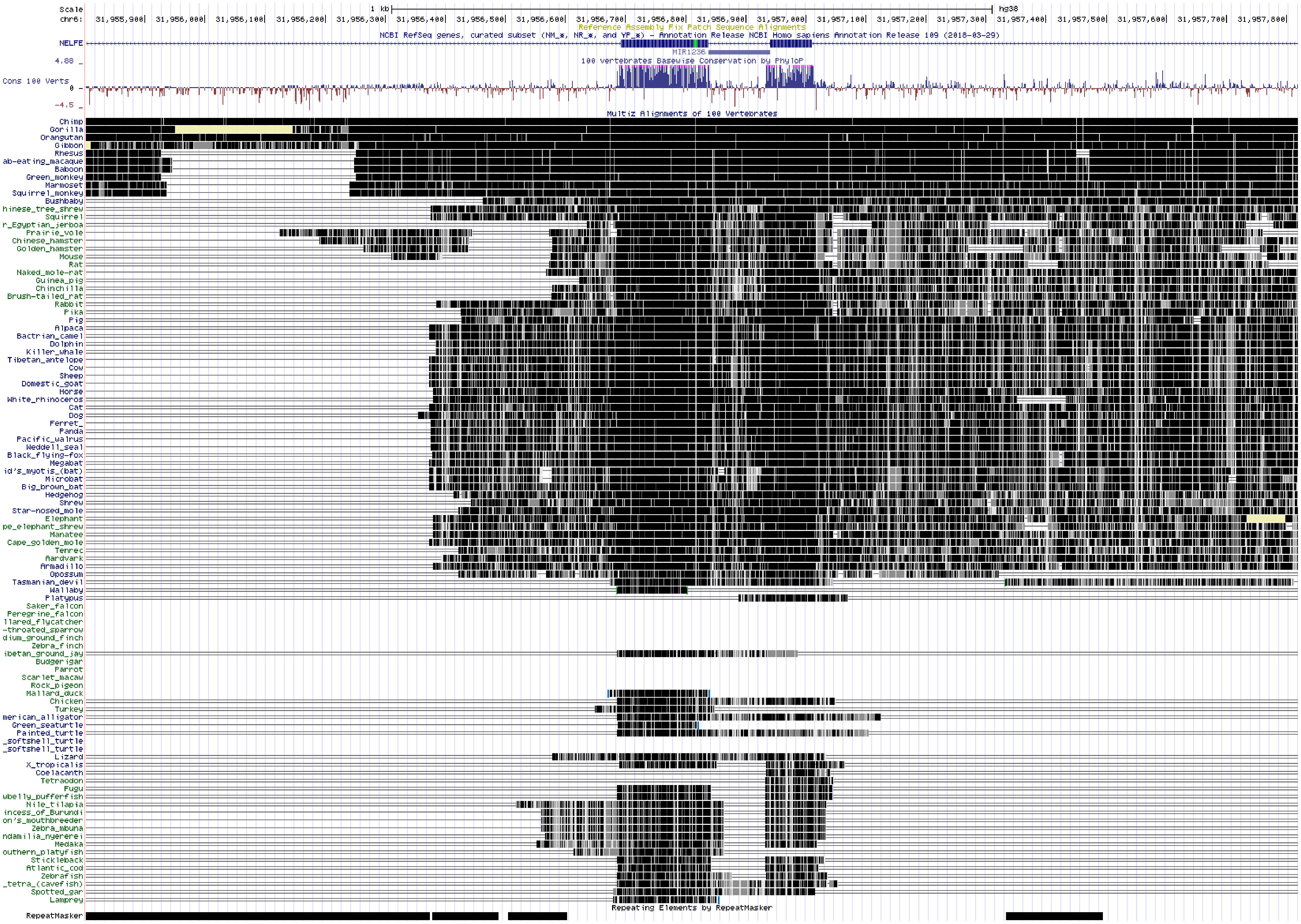


**Appendix Figure S7. Conservation of the miR-1236 sequence among vertebrate animals.**

The genomic sequences (including miR-1236 and the surrounding exons and introns derived from several vertebrate animals) were aligned, and their sequence homologies were compared using the UCSC genome browser (http://genome.ucsc.edu).

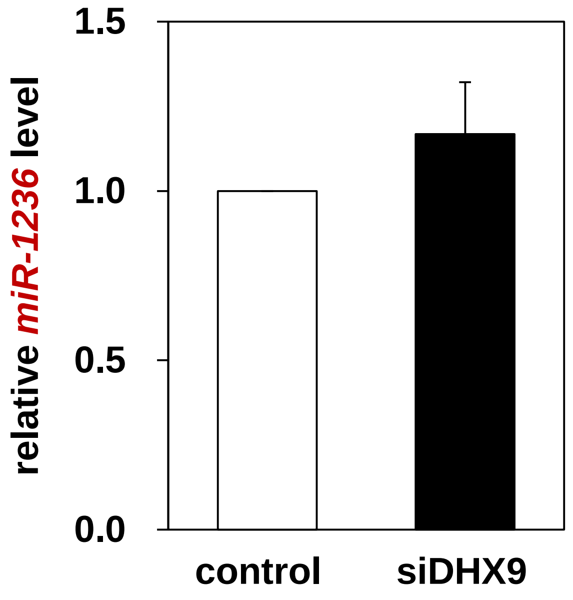


**Appendix Figure S8. Depleting DHX9 did not cause gross changes in the amount of miR-1236 associated with AGO3.**

Following transfection of KGN cells with control siRNA or siDHX9, AGO3-mediated RISC-associated RNAs were isolated by immunoprecipitation using an anti-AGO3 antibody. The RNA was extracted using an acidic phenol: chloroform mixture (5: 1, pH 4.3) and precipitated with isopropanol in the presence of 10% of 3M NaOAc (pH 5.2). The relative abundance of miR-1236 was validated using TaqMan® microRNA assays. The data (mean ± SEM) are from three independent experiments, performed in triplicate.
