## Supplement Table for "An alternative miRISC targeting a coding mutation site in *FOXL2* links to granulosa cell tumor"

**Appendix Table S1. Predicted miRNAs with a possible seed match around 402C>G site of *FOXL2***

| miRNAs | C402G | | WT | | position[c] | alignment[d] |
|---|---|---|---|---|---|---|
| | ΔΔG[a] | Seed[b] | ΔΔG[a] | Seed[b] | | |
| hsa-miR-1236 | -9.26 | 8:1:0 | -9.16 | 6:1:0 | 401 | 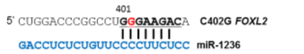 |
| hsa-miR-145 | -2.14 | 6:1:1 | N/A | N/A | 398 | 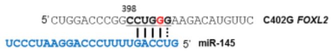 |
| hsa-miR-204 | -10.9 | 6:1:0 | -4.3 | 6:1:0 | 418 | 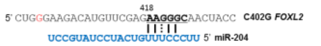 |
| hsa-miR-942 | -4.59 | 8:1:1 | -1.39 | 6:1:0 | 399 | 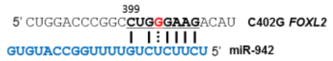 |
| hsa-miR-1266 | -6.63 | 8:1:1 | -3.1 | 6:1:0 | 396 | 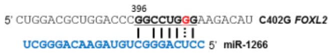 |
| hsa-miR-632 | -2.42 | 7:1:0 | -0.91 | 7:1:0 | 403 | 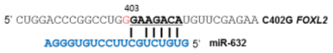 |
| hsa-miR-484 | -10.97 | 6:0:1 | -10.04 | 6:0:1 | 396 | 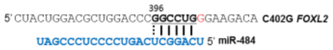 |
| hsa-miR-940 | -14.74 | 8:1:0 | -14.3 | 8:1:0 | 393 | 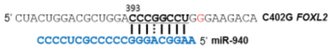 |
| hsa-miR-1295 | -7.46 | 8:1:1 | -7.13 | 8:1:1 | 394 | 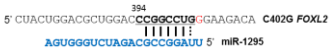 |
| hsa-miR-423-3p | -9.94 | 8:1:1 | -9.67 | 8:1:1 | 391 | 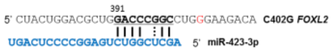 |
| hsa-miR-1287 | -8.69 | 8:1:1 | -8.42 | 8:1:1 | 391 | 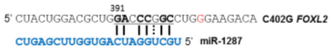 |
| hsa-miR-661 | -14.99 | 8:1:1 | -14.9 | 8:1:1 | 392 | 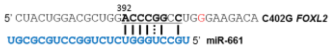 |
| hsa-miR-485-5p | -5.66 | 8:1:1 | -5.63 | 8:1:1 | 394 | 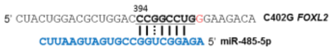 |
| hsa-miR-127-3p | -6.69 | 8:1:1 | -6.71 | 8:1:1 | 389 | 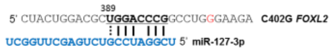 |
| hsa-miR-149 | -9.96 | 8:1:1 | -10.04 | 8:1:1 | 390 | 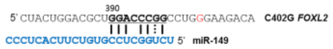 |
| hsa-miR-671-3p | -6.74 | 8:1:1 | -6.82 | 8:1:1 | 390 | 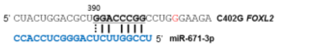 |
| hsa-miR-198 | -3.13 | 8:1:0 | -4.07 | 8:1:0 | 386 | 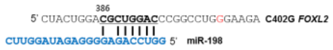 |
| hsa-miR-601 | -2.6 | 8:1:1 | -3.98 | 8:1:1 | 387 | 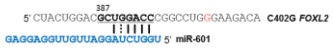 |

| | | | | | | |
|---|---|---|---|---|---|---|
| hsa-miR-187 | -8.11 | 8:0:1 | -11.81 | 8:0:1 | 404 | 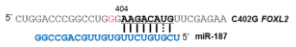  |

[a]ΔΔG = ΔGduplex – ΔGopen, (kcal/mol). (ΔGduplex, the energy gained by binding of the miRNA to the target, ΔGopen, the energy required to make the target region accessible for miRNA binding) The lower (more negative) its value, the stronger the binding of the microRNA to the given site is expected to be. [b]"X:Y:Z" in the seed column, the size of the seed (X), the number of mismatches (Y) and the number of G:U wobble pairs (Z). Those with the size of seed more than 5 were screened. [c]position, the first position of the seed sequence. [d]alignment, sequence alignment of miRNAs with the putative binding sites in the 402C>G *FOXL2* mRNA. N/A, not available.
